# CAP1 and cofilin1 cooperate in neuronal actin dynamics, growth cone function and neuron connectivity

**DOI:** 10.1101/2020.08.12.247932

**Authors:** Felix Schneider, Thuy-An Duong, Isabell Metz, Jannik Winkelmeier, Christian A. Hübner, Ulrike Endesfelder, Marco B. Rust

## Abstract

Neuron connectivity depends on growth cones that navigate axons through the developing brain. Growth cones protrude and retract actin-rich structures to sense guidance cues. These cues control local actin dynamics and steer growth cones towards attractants and away from repellents, thereby directing axon outgrowth. Hence, actin binding proteins (ABPs) moved into the focus as critical regulators of neuron connectivity. We found cyclase-associated protein 1 (CAP1), an ABP with unknown brain function, abundant in growth cones. Super-resolution microscopy and live cell imaging combined with pharmacological approaches on hippocampal neurons from gene-targeted mice revealed a crucial role for CAP1 in actin dynamics that is critical for growth cone morphology and function. Growth cone defects in mutant neurons compromised neuron differentiation and was associated with impaired neuron connectivity in CAP1 mutant brains. Mechanistically, we found that CAP1 and cofilin1 synergistically control growth cone actin dynamic and morphology. Together, we identified CAP1 as a novel actin regulator in growth cone that is relevant for neuron connectivity.

## Introduction

The formation of complex brain circuits depends on directed extension of axonal projections from neurons to their specific targets. During development, axons are guided by a ‘hand-shaped’ structure at their distal tip termed growth cone (Ramón y Cajal, 1909). The peripheral domain of growth cones, the leading edge, is highly enriched in actin filaments (F-actin) and contains receptors that translate guidance cues into intracellular signaling activities that act upstream of actin-binding proteins (ABPs) to control F-actin assembly and disassembly. Local actin dynamics steer the growth cone towards attractive and away from repellent cues and, hence, navigate axon through the developing brain (Gomez and Letourneau, 2014). To fulfill its sensory function and to explore its environment, the leading edge is highly motile and persistently protrudes and retracts actin-rich structures such as finger-like filopodia or sheet-like lamellipodia (Dent et al., 2011). Hence, actin dynamics play a central role in growth cone function, and ABPs moved into the focus as critical regulators of neuron connectivity and brain development (Gomez and Letourneau, 2014; Omotade et al., 2017; Vitriol and Zheng, 2012).

Actin dynamics are controlled by ABPs with different biochemical functions including actin nucleation and polymerization, F-actin depolymerization and severing, nucleotide exchange on globular actin monomers (G-actin), and blocking of barbed or pointed ends (Pollard, 2017). Although numerous ABPs including the actin depolymerizing protein cofilin1 have been implicated in growth cone motility, our knowledge about actin regulatory mechanisms in growth cones is still fragmented (Dent et al., 2011; Gomez and Letourneau, 2014; Omotade et al., 2017; Vitriol and Zheng, 2012). By exploiting recombinant proteins and yeast mutant strains, recent studies implicated cyclase-associated protein (CAP) in both F-actin disassembly and nucleotide exchange on G-actin (Johnston et al., 2015; Kotila et al., 2018; Kotila et al., 2019; Shekhar et al., 2019). Unlike yeast, mammals possess two CAP family members, CAP1 and CAP2, with different expression patterns. CAP2 is abundant in striated muscles and brain (Bertling et al., 2004), and recent mouse studies implicated CAP2 in heart physiology, skeletal muscle development and synaptic function (Peche et al., 2012; Field et al., 2015; Kepser et al., 2019; Pelucchi et al., 2020). Instead, CAP1 expression is less restricted, and its physiological functions remained unknown, also because appropriate animal models were lacking (Jang et al., 2019).

We found CAP1 expression during neuron differentiation and abundance in growth cones and therefore hypothesized a crucial role in growth cone function and brain development. To test our hypothesis, we generated a conditional mouse model and inactivated CAP1 during brain development. Analyses of hippocampal neurons from mutant mice and brain histology revealed a role for CAP1 in neuron differentiation and neuron connectivity. Further, we identified CAP1 as a novel regulator of actin dynamics that controls growth cone motility and morphology in response to guidance cues. Rescue experiments in neurons lacking either CAP1 or CAP1 together with cofilin1 revealed that CAP1 and cofilin1 synergistically control neuronal actin dynamics and growth cone morphology.

## Results

### CAP1 is relevant for neuron connectivity in the mouse brain

Immunoblots revealed presence of CAP1 in mouse cerebral cortex lysates throughout embryonic development, between embryonic day (E) 12.5 and postnatal day (P) 0 (Fig. 1A). At P0, CAP1 expression levels in hippocampal lysates were similar to those in cerebral cortex lysates (Fig. 1B). We therefore hypothesized a function for CAP1 during brain development. Systemic CAP1 mutant mice died during embryonic development and, hence, were not useful to test our hypothesis (Jang et al., 2019). We therefore generated mice carrying a targeted CAP1 allele with exon 3 being flanked by loxP sites (CAP1^flx/flx^; Fig. 1C). Brain-specific CAP1 inactivation was achieved by crossing CAP1^flx/flx^ mice with Nestin-Cre transgenic mice (Tronche et al., 1999). Compared to CAP1^flx/flx^ littermates (CTR), CAP1 mRNA levels were strongly reduced in total brain lysates from E18.5 CAP1^flx/flx,Nestin-Cre^ mice termed CAP1-KO (Fig. 1D; CTR: 1.00±0.15, CAP1-KO: 0.17±0.06, n=3, P<0.05), and CAP1 protein levels dropped below detection limit (Fig. 1E). Hence, CAP1-KO mice were valuable tools to study the brain function of CAP1.

**Figure 1.**
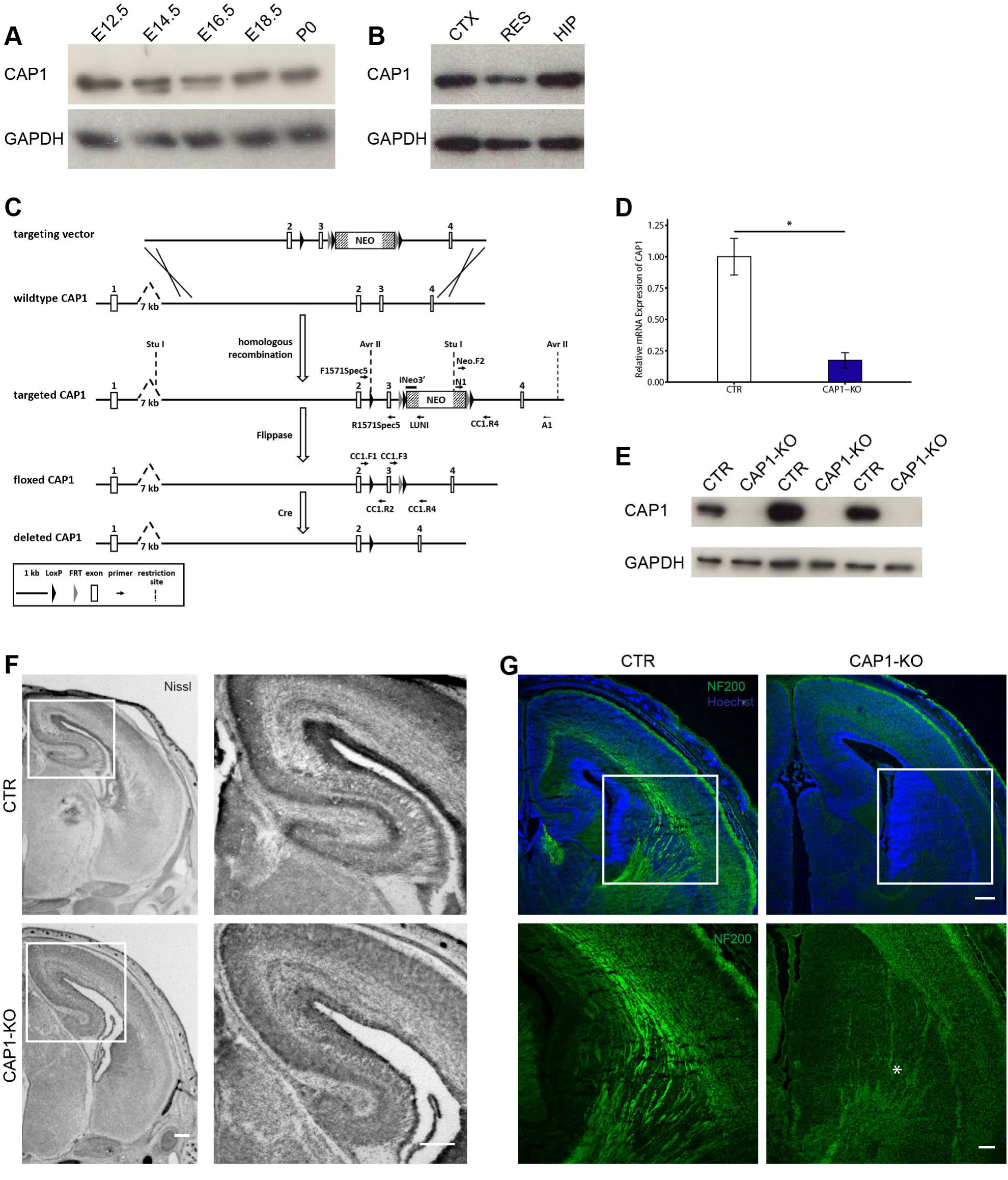
Reduced neuron connectivity in CAP1 mutant mouse brain. **(A)** Immunoblots showing presence of CAP1 in cerebral cortex lysates during embryonic development, from embryonic day (E) 12.5 to postnatal day (P) 0. GAPDH was used as loading control. **(B)** Immunoblots showing equal CAP1 expression levels in cerebral cortex (CTX) and hippocampus (HIP) lysates at P0. Additionally, lysates of residual brains (RES) were probed. GAPDH was used as loading control. **(C)** Scheme showing targeting strategy for CAP1. The targeted CAP1 allele contained two loxP sites flanking exon 3 and a neomycin resistance cassette flanked by Frt sites. Upon successful homologous recombination in ES cells and blastocyst injection, the neomycin resistance cassette was removed by crossing gene targeted mice with a flippase deleter mouse. CAP1 deletion during brain development was achieved by crossing CAP1^flx/flx^ mice with CAP1^+/flx,Nestin-Cre^ mice. **(D)** Quantitative PCR (qPCR) showing reduced CAP1 mRNA levels in brain lysates from E18.5 CAP1-KO mice. **(E)** Immunoblots (three biological replicates) showing absence of CAP1 from E18.5 CAP1-KO brains. GAPDH was used as a loading control. **(F)** Nissl staining of transversal brain sections from E18.5 CTR and CAP1-KO mice. Boxes indicate areas shown at higher magnification. **(G)** Antibody staining of transversal brain sections from E18.5 CTR and CAP1-KO mice against the axonal marker NF200 (green). Boxes indicate areas shown at higher magnification. Asterisk indicates missing axon fibers in CAP1-KO brain. Sections were counterstained with the DNA dye Hoechst (blue). Scale bars (in µm): 250 (F, G). *: P<0.05.

When crossing CAP1^flx/flx^ with CAP1^+/flx,Nestin-Cre^ mice, we found the expected 25% CAP1-KO mice among all offspring (n>300) at E18.5. However, CAP1-KO mice died shortly after birth, thereby restricting our histological analysis to embryonic stages. Antibody staining against the mitotic marker phospho-Histone 3 or the neural stem cell markers Pax6 and Tbr2 revealed no differences between CTR and CAP1-KO mice (data not shown). In line with this, Nissl staining as well as antibody staining against the neuron layer markers Tbr1 and Ctip2 revealed no obvious defects in cerebral cortex or hippocampus anatomy at E18.5 (Figs. 1F, S1). Conversely, antibody staining against the axon marker neurofilament revealed clearly less fiber tracks in CAP1-KO brains, e.g. in the cortical intermediate zone or in the striatum (Fig. 1G). Hence, neuron connectivity was compromised in CAP1-KO mice.

### CAP1 inactivation impaired neuron differentiation

Compromised neuron connectivity in CAP1 mutant brains could be caused by impaired neuron differentiation. To test a role for CAP1 in neuron differentiation, we chose hippocampal neurons isolated from E18.5 mice as a cellular system, which expressed substantial CAP1 levels when kept *in vitro* (Fig. 2A). We stained neurons with an antibody against the neurite marker doublecortin (Dcx, Fig. 2B), which allowed us to categorized neurons according to their differentiation stage (Dotti et al., 1988). We counterstained neurons with fluorescent phalloidin that labels F-actin (Melak et al., 2017). After five hours *in vitro* (HIV5), we found the majority of CTR and CAP1-KO neurons in stage 1, i.e. they formed lamellipodia, but not yet neurites (Fig. 2C; (in %) CTR: 69.23±3.90; CAP1-KO 84.63±2.76, n>300 neurons from 3 mice). All other neurons possessed minor neurites, but not yet an axon, and were assigned to stage 2 (CTR: 30.77±3.90, CAP1-KO: 15.37±2.76). Compared to CTR, the fraction of stage 2 neurons was halved in CAP1-KO cultures, and the stage distribution was different between both groups (P<0.001). Stage distribution differences were even more pronounced at later time points. After one day *in vitro* (DIV1), a minority of CTR neurons remained in stage 1, while the vast majority has been in stage 2 and some neurons already possessed an axon and reached stage 3 ((in %) stage 1: 10.21±1.79, stage 2: 81.64±3.17, stage 3: 8.15±2.27; n>300/3). Conversely, approximately half of all CAP1-KO neurons remained in stage 1 and virtually no stage 3 CAP1-KO neurons were present ((in %) stage 1: 51.31±2.73, stage 2: 48.38±2.57, stage 3: 0.31±0.31; n>300/3; P<0.001). At DIV2, the fraction of stage 3 CTR neurons increased to roughly one third, and almost all other neurons were in stage 2 ((in %) stage 1: 4.33±1.67, stage 2: 59.76±3.85, stage 3: 35.91±3.47; n>230/3). Instead, only a few CAP1-KO neurons reached stage 3, while one third remained in stage 1 ((in %) stage 1: 31.85±4.16, stage 2: 64.80±4.32, stage 3: 3.35±1.21; n>230/3, P<0.001). Antibody staining against the axon marker tau-1 in DIV2 cultures confirmed presence of axons in stage 3 CTR and CAP1-KO neurons, thereby proving correct categorization (Fig. S2A). Notably, treatment of isolated neurons with 1 µM cytochalasin D (CYTOD), a mycotoxin that prevents polymerization of G-actin (Flynn et al., 2012), normalized stage distribution of CAP1-KO neurons at DIV2 (Fig. 2D-E; (in%) CTR-DMSO: stage 1: 7.08±1.23, stage 2: 70.16±1.84, stage 3: 22.77±1.80; CAP1-KO-DMSO: stage 1: 28.92±2.26, stage 2: 62.55±1.80, stage 3: 8.53±2.33, P<0.001, n>200/3; CAP1-KO-CYTOD: stage 1: 4.17±1.63, stage 2: 74.72±3.89, stage 3: 21.11±3.53, n>220/3). Upon CYTOD treatment, stage distribution of CAP1-KO neurons was different from DMSO-treated CAP1-KO neurons (P<0.001), but not from DMSO-treated CTR neurons (P=0.69). Together, our data revealed a role for CAP1 in neuron differentiation and suggested that it controls neuron differentiation via an actin-dependent mechanism.

**Figure 2.**
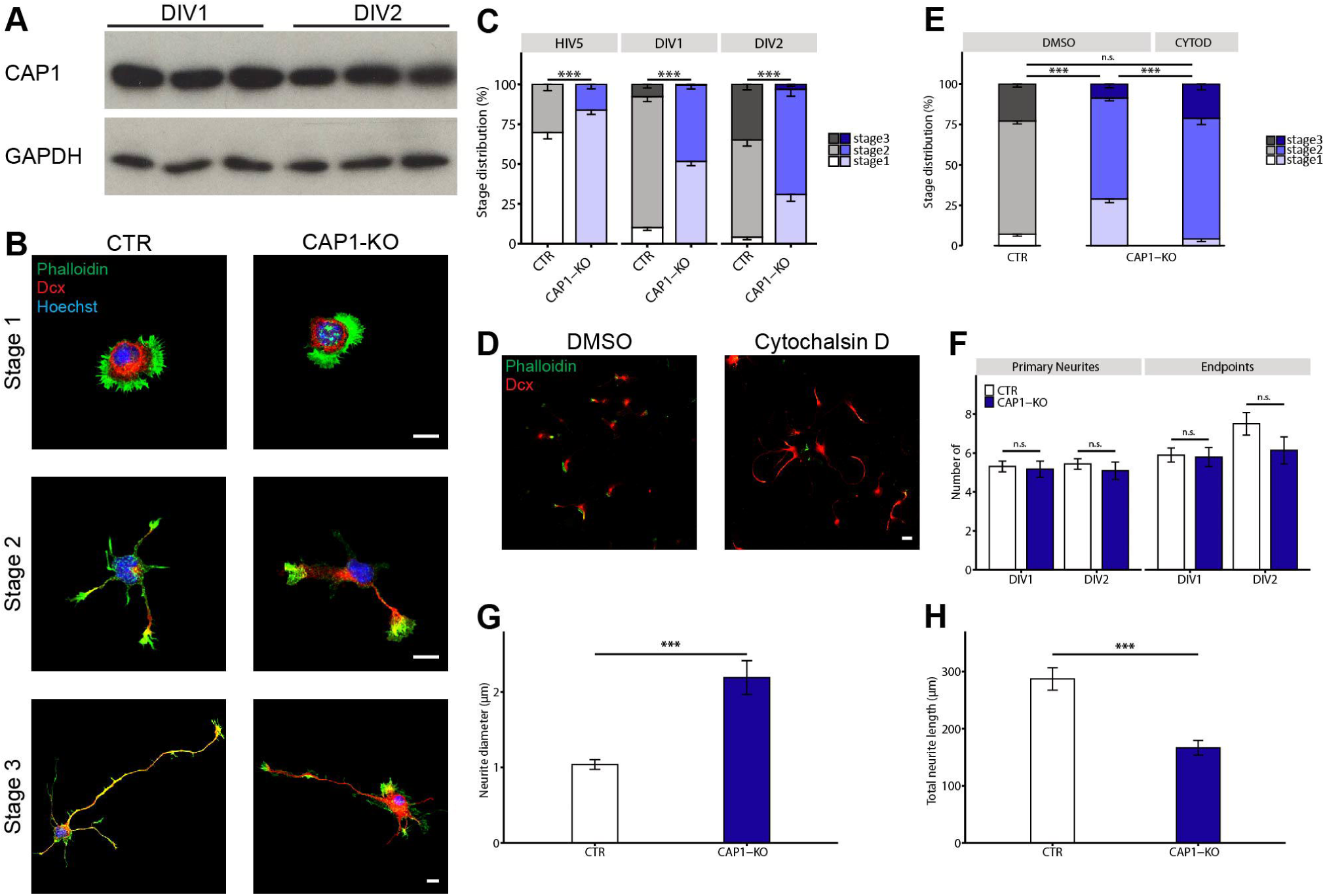
Impaired differentiation and altered neurite morphology in CAP1-KO neurons. **(A)** Immunoblots (three biological replicates) showing presence of CAP1 in lysates from cultured hippocampal neurons at DIV1 and DIV2. GAPDH was used as loading control. **(B)** Representative hippocampal neurons from CTR and CAP1-KO mice at differentiation stages 1, 2 and 3 (Dotti et al., 1988). Neurons were stained with an antibody against the neuronal marker doublecortin (Dcx, red), the F-actin marker phalloidin (green) and Hoechst (blue). **(C)** Stage distribution for CTR and CAP1-KO neurons at HIV5, DIV1 and DIV2. **(D)** Representative hippocampal CAP1-KO neurons treated either with DMSO or cytochalasin D (CYTOD). Neurons were stained with an antibody against Dcx (red) and phalloidin (green). **(E)** Stage distribution of DMSO-treated CTR and CAP1-KO neurons and CYTOD-treated CAP1-KO neurons. **(F)** Number of primary neurites and neurite endpoints in stage 2 CTR and CAP1-KO neurons. **(G)** Neurite width in stage 2 neurons. **(H)** Neurite length in stage 2 neurons. Scale bar (in µm): 10 (B), 25 (D); n.s.: P≥0.05, ***: P<0.001.

### CAP1 inactivation altered neurite length and width

Next, we determined neuron morphology by counting the numbers of primary neurites and neurite endpoints. Both parameters were unchanged in stage 2 or stage 3 CAP1-KO neurons (Figs. 2F, S2B; DIV1: stage 2: neurites: CTR: 5.31±0.28, CAP1-KO: 5.17±0.41, P=0.33; endpoints: CTR: 5.90±0.36, CAP1-KO: 5.79±0.49, P=0.52, n=30/3; DIV2: stage 2: neurites: CTR: 5.44±0.27, CAP1-KO: 5.09±0.45, P=0.37; endpoints: CTR: 7.50±0.58, CAP1-KO: 6.14±0.69, P=0.08, n=20/3; stage 3: neurites: CTR: 5.21±0.26, CAP1-KO: 5.30±0.62, P=0.90; endpoints: CTR: 7.71±0.85, CAP1-KO: 6.90±0.99, P=0.64, n=10/6). Further, by exploiting Dcx-labelled neurons (Fig. 2B), we determined neurite length and width. While neurite width was doubled in CAP1-KO neurons (Fig. 2G; (in µm) CTR: 1.04±0.06, CAP1-KO: 2.19±0.22, P<0.001), total neurite length was reduced by 40% (Fig. 2H; (in µm) CTR: 286.99±19.59, CAP1-KO: 166.58±12.79, P<0.001). Together, neurite numbers and branching were unchanged in CAP1-KO neurons, but their neurites were thicker and shorter.

### CAP1 inactivation impaired growth cone morphology and motility

To decipher the CAP1-dependent mechanism during neuron differentiation, we determined its subcellular localization. Antibody staining revealed CAP1 abundance in growth cones (Figs. 3A), which we identified by phalloidin labelling of F-actin. Line scans showed overlapping fluorescent intensity profiles for CAP1 and phalloidin (Fig. 3B). Localization in growth cones was confirmed by overexpression of green fluorescence protein (GFP)-tagged CAP1, which unlike GFP in control neurons, was enriched in growth cones (Fig. S3). Hence, during neuron differentiation CAP1 was enriched in growth cones.

**Figure 3.**
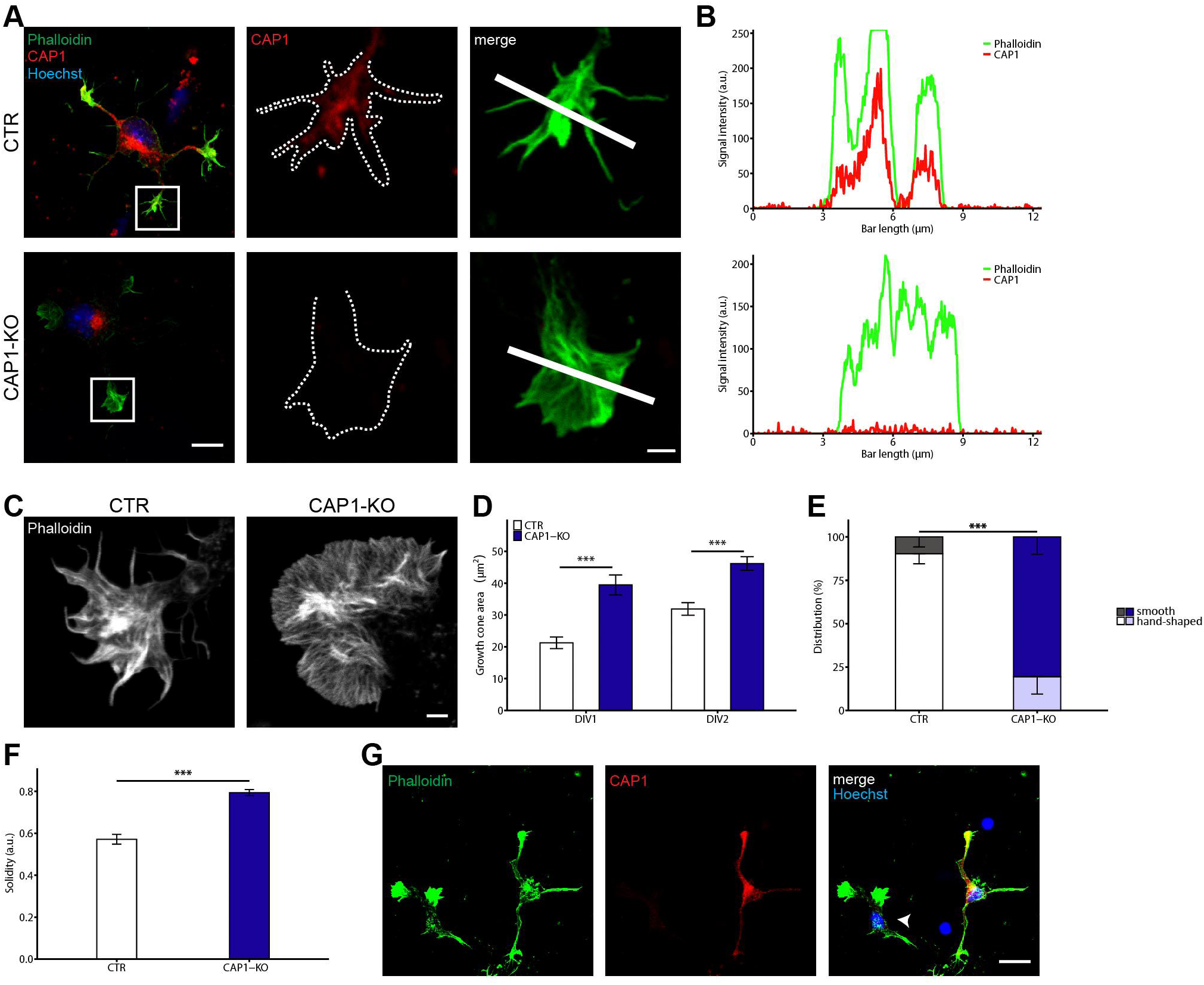
Increased size and altered morphology of growth cones from CAP1-KO neurons. **(A)** Antibody staining against CAP1 (red) in CTR and CAP1-KO neurons. Neurons were counterstained with phalloidin (green) and Hoechst (blue). Boxes indicate areas shown at higher magnification, dashed lines outline growth cones. **(B)** CAP1 and phalloidin fluorescence intensity profiles along white lines in A. **(C)** Representative micrographs of phalloidin-stained growth cones from stage 2 neurons. **(D)** Growth cone size in CTR and CAP1-KO neurons at DIV1 and DIV2. **(E)** Growth cone fraction from CTR and CAP1-KO neurons with ‘hand-shaped’ or smooth morphology. **(F)** Growth cone shape index solidity for CTR and CAP1-KO neurons. **(G)** Mixed cultures of CTR and CAP1-KO neurons stained with an antibody against CAP1 (red) and counterstained with phalloidin (green) and Hoechst (blue). Arrowhead marks a CAP1-KO neuron. Scale bars (in µm): 2 (A high magnification, C), 10 (A low magnification), 25 (G). ***: P<0.001.

To test whether CAP1 is functionally relevant in growth cones, we determined their size in stage 2 neurons. Compared to CTR, growth cones size was increased by 85% and 45 % in CAP1-KO neurons at DIV1 and DIV2, respectively (Fig. 3C-D; (in µm^2^) DIV1: CTR: 21.26±1.83, CAP1-KO: 39.46±3.13, n=180/6, P<0.001; DIV2: CTR: 31.90±1.98, CAP1-KO: 46.19±2.16, n=90/3, P<0.001). Further, we noted an altered growth cone morphology in CAP1-KO neurons. Opposite to CTR neurons, only a minority of 20% displayed the typical ‘hand-shaped’ morphology, while most CAP1-KO growth cones appeared rather smooth (Fig. 3E; (in %) hand-shaped: CTR: 90.30±5.78, CAP1-KO: 19.44±10.02; smooth: CTR: 9.70±5.78, CAP1-KO: 80.55±10.02, P<0.001, n>25/3). A smooth growth cone morphology in CAP1-KO neurons was also evident from a 40% increase in shape index solidity (Fig. 3F; CTR: 0.57±0.02, CAP1-KO: 0.79±0.01, P<0.001, n=26/3). Similar changes in growth cone morphology were present in CAP1-KO neurons co-cultured with CTR neurons (Fig. 3G). Together, CAP1 controls growth cone size and morphology, suggesting a role in growth cone function. Indeed, growth cones from CAP1-KO neurons were clearly less motile when compared to CTR neurons (Movies S1, S2). Since F-actin constitutes the major structural backbone relevant for growth cone morphology and motility (Gomez and Letourneau, 2014; Omotade et al., 2017), we hypothesized a CAP1 function in regulating growth cone actin cytoskeleton.

### CAP1 controls F-actin organization and dynamics in growth cones

To test this hypothesis, we first exploited phalloidin-labelled growth cones and examined F-actin organization by direct stochastic optical reconstruction microscopy (dSTORM; Figs. 4A, S4). This approach revealed an overall normal F-actin density in CAP1-KO growth cones (Fig. 4B; (in localization numbers/µm^2^) CTR: 5,519±562; CAP1-KO: 5,115±525, n=18/5, P=0.53). However, CAP1-KO growth cones displayed two obvious alterations: the number of filopodia was reduced by 75% (Fig. 4C; CTR: 9.26±0.72; CAP1-KO: 2.18±.0.35, n=18/5, P<0.001), and most CAP1-KO growth cones lack a clearly defined central domain, which normally contains only few F-actin structures (Dupraz et al., 2019; Lowery and Van Vactor, 2009). Hence, dSTORM revealed a normal density, but an altered F-actin organization in CAP1-KO growth cones.

**Figure 4.**
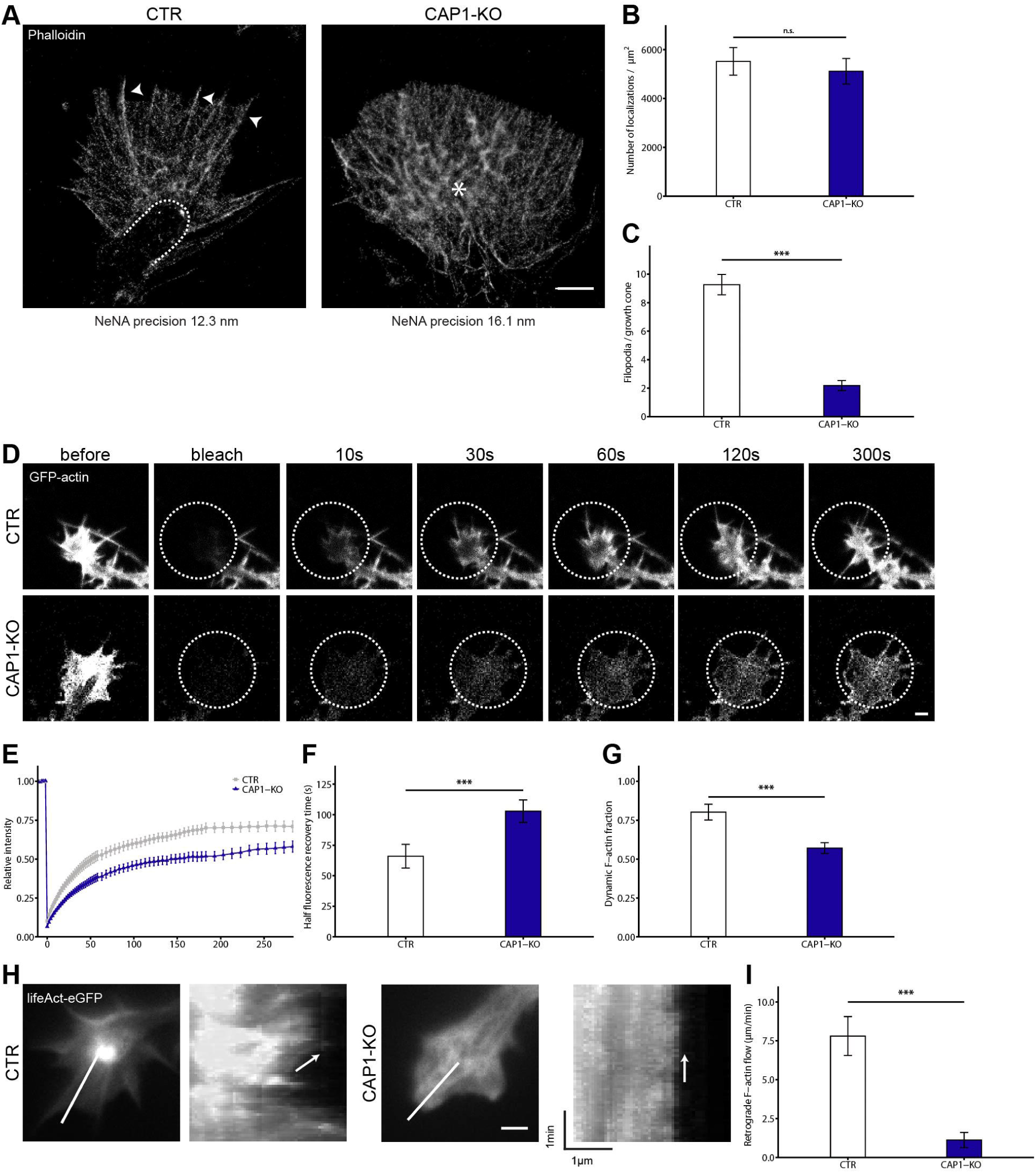
CAP1 controls F-actin organization and dynamics in growth cones. **(A)** Representative micrographs of phalloidin-stained growth cones from CTR and CAP1-KO neurons acquired by dSTORM. Dashed line marks border between C domain and T zone. Arrowheads exemplarily point to filopodia. Asterisk indicates lack of C domain in CAP1-KO growth cone. **(B)** F-actin density (phalloidin localizations normalized to growth cone size) in CTR and CAP1-KO growth cones. **(C)** Filopodia number per growth cone in CTR and CAP1-KO neurons. **(D)** Image sequence of GFP-actin in CTR and CAP1-KO growth cones during FRAP experiments. Dashed lines encircle bleached areas. **(E)** Recovery curves for GFP-actin in growth cones from CTR and CAP1-KO neurons. **(F)** Half-recovery time for GFP-actin in growth cones from CTR and CAP1-KO neurons. **(G)** Dynamic actin fraction in growth cones from CTR and CAP1-KO neurons. **(H)** Representative micrographs of growth cones from LifeAct-GFP-expressing CTR and CAP1-KO neurons. Lines indicate where kymographs (shown on the right) have been generated from. Arrows indicate the retrograde F-actin flow. **(I)** Velocity of retrograde F-actin flow in growth cones from CTR and CAP1-KO neurons. Scale bars (in µm): 2 (A, D, H); n.s.: P≥0.05, ***: P<0.001.

Further, we examined whether actin dynamics was altered in CAP1-KO growth cones. We therefore performed fluorescence recovery after photobleaching (FRAP) experiments in GFP-actin-expressing neurons to determine actin turnover within a time frame of 300 s upon bleaching (Movies S3, S4). In CTR growth cones, GFP-actin rapidly recovered with a mean half-recovery time (t_½_) of 65.99±9.72 s (Fig. 4D-F). Fluorescence recovery was much slower in CAP1-KO growth cones as indicated by a roughly 60% increase in t_½_ (102.88±9.17 s, n=40/3, P<0.001). Moreover, the dynamic actin fraction that recovered within 300 s was reduced by 30% in CAP1-KO growth cones (Fig. 4G; CTR: 0.80±0.05, CAP1-KO: 0.57±0.03, P<0.001). We also expressed LifeAct-GFP to visualize F-actin in living neurons (Riedl et al., 2008). Compared to CTR neurons, F-actin appeared less dynamic in growth cones from CAP1-KO neurons (Movies S5, S6). Kymograph analysis allowed us to determine retrograde F-actin flow and, hence, to quantify F-actin dynamics in growth cones (Flynn et al., 2012). In CTR growth cones, the average retrograde flow velocity was 7.81±1.25 µm/min (Fig. 4H-I), while in CAP1-KO neurons the velocity was seven-fold reduced (1.12±0.49 µm/min, n=25/3, P<0.001). Together, our data demonstrated a crucial role for CAP1 in growth cone F-actin organization and dynamics.

### CAP1 controls growth cone size downstream of guidance cues

Guidance cues act upstream of growth cone actin dynamics and thereby control neurite outgrowth and neuron connectivity. Next, we tested whether CAP1 was relevant for guidance cue-induced morphological changes in growth cones. We therefore determined growth cones upon treatment with either the attractant brain-derived neurotrophic factor (BDNF) or the repellent cues Ephrin A5 (EphA5), Semaphorin 4D (Sema4D) and Slit-1. Compared to PBS-treated neurons, growth cones were enlarged by 60% in CTR neurons upon BDNF treatment (Fig. 5A-B; (in µm^2^) PBS: 33.09±2.08, BDNF: 52.02±2.68, P<0.001, n=80/3). Instead, BDNF did not enlarge growth cones in CAP1-KO neurons ((in µm^2^) PBS: 64.24±3.70, BDNF: 58.33±3.03, P=0.2, n=60/3), and growth cone size was similar in CTR and CAP1-KO neurons upon BDNF treatment (P=0.20). Hence, CAP1-KO growth cones failed in responding to the attractant cue BDNF.

**Figure 5.**
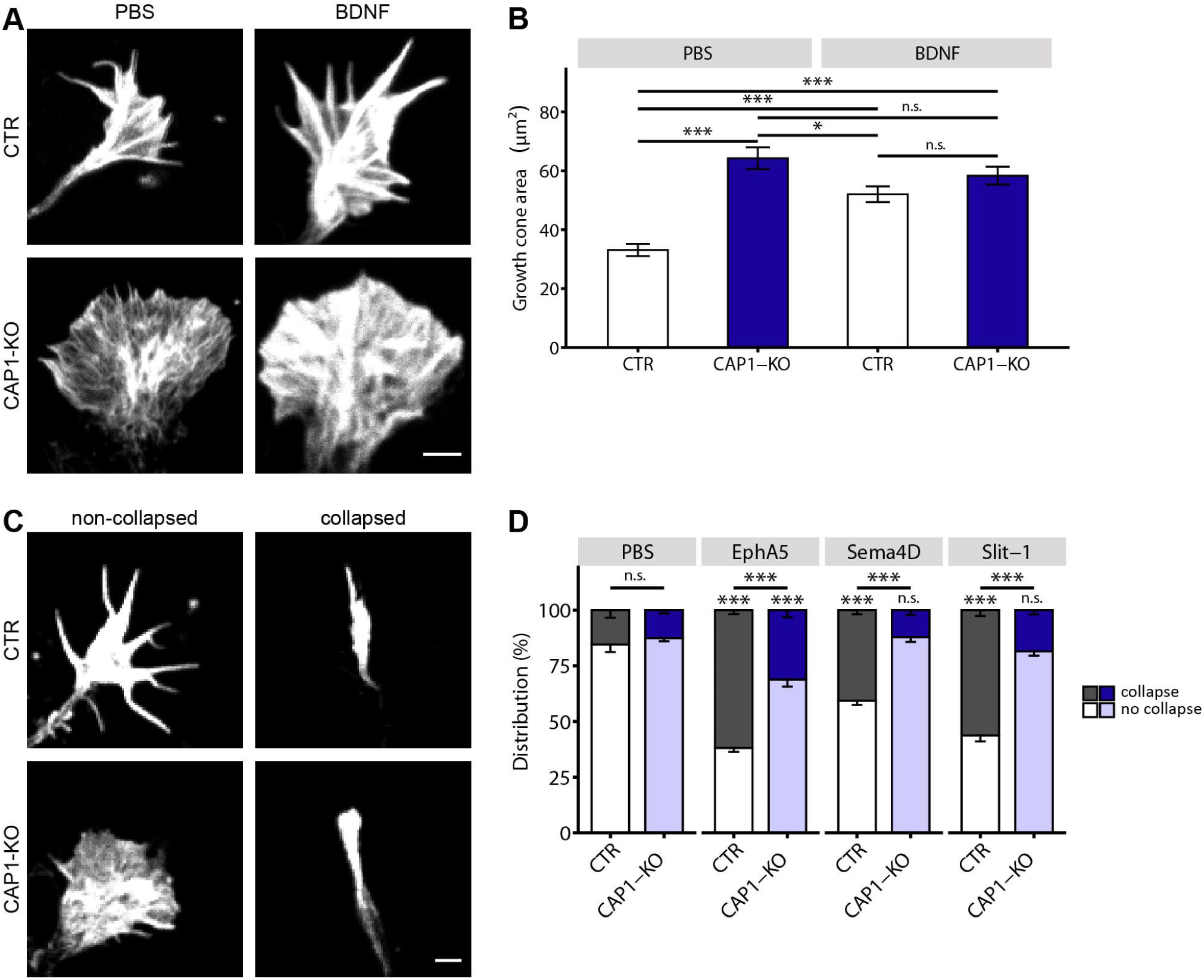
CAP1 acts downstream of guidance cues in growth cones. **(A)** Phalloidin-labeled growth cones from CTR and CAP1-KO neurons treated with either PBS or BDNF. **(B)** Growth cone size in CTR and CAP1-KO neurons treated with either PBS or BDNF. **(C)** Phalloidin-stained non-collapsed and collapsed growth cones from CTR and CAP1-KO neurons. **(D)** Fraction of collapsed and non-collapsed growth cones from CTR and CAP1-KO neurons treated with either PBS or the repellent cues EphrinA5 (EphA5), Semaphorin D (Sema4D) or Slit-1. Comparisons between PBS-treated control conditions and treatment with either EphA5, Sema4D or Slit-1 are indicated by asterisks directly above bars. Scale bars (in µm): 2 (A, C); n.s.: P≥0.05, *: P<0.05, ***: P<0.001.

CAP1-KO neurons were also impaired in growth cone collapse downstream of the repellent cues EphA5, Sema4D and Slit-1. EphA5 fourfold increased the fraction of collapsed growth cones in CTR neurons (Fig. 5C-D; (in %) PBS: 15.43±3.41, EphA5: 61.87±1.79, n=200/3, P<0.001). EphA5 also increased the fraction of collapsed CAP1-KO growth cones ((in %) PBS: 12.52±1.47, EphA5: 31.13±3.20, n=200/3, P<0.001), but the EphA5 effect was smaller compared to CTR neurons (P<0.001). Sema4D and Slit-1 both induced growth cone collapse in CTR neurons and increased the collapsed growth cone fraction by a factor of 2.6 and 3.7, respectively (Sema4D: 40.70±1.85, P<0.001; Slit-1: 56.29±2.64, P<0.001). Conversely, neither Sema4D nor Slit-1 increased the collapsed growth cone fraction in CAP1-KO neurons (Sema4D: 12.12±2.13, P=0.85; Slit-1: 18.50±1.93, P=0.42). Together, CAP1 acts downstream of BDNF and the repellent cues EphA5, Sema4D and Slit-1, thereby implicating CAP1 in signaling cascades that are relevant for directed neurite outgrowth and neuron connectivity in the mammalian brain (Lowery and Van Vactor, 2009).

### The helical folded domain is critical for CAP1 function in growth cones

CAP1 is a multidomain protein comprising several conserved domains capable of actin or ABP binding (Fig. 6A). By expressing CAP1 mutants in CAP1-KO neurons, we next tested which domain was relevant in growth cones. In these experiments, we used growth cone size as a readout, which showed the most robust difference between CTR and CAP1-KO (Fig. 3). In control experiments, we expressed GFP-tagged wild-type (WT) CAP1, which did not change growth cone size in CTR, but normalized size in CAP1-KO neurons (Fig. 6B-C; (in µm^2^): CTR: GFP: 22.71±1.52, WT-CAP1: 25.90±2.16, n=100/3, P=0.38; CAP1-KO: GFP: 42.02±2.55, WT-CAP1: 27.54±2.43, n=80/3, P<0.001). Unlike WT-CAP1, deletion constructs lacking either the N-terminal 213 amino acid (aa) residues (CAP1-Δ1-213) or the last 156 residues (CAP1-Δ319-474) did not normalize growth cone size in CAP1-KO neurons (Δ1-213: 35.64±2.74, n=80/3, P=0.09; Δ319-474: 38.92±2.34, n=70/3, P=0.46). To narrow down CAP1 domains relevant in growth cones, we mutated CAP1’s helical folded domain (HFD), ii) proline-rich stretch (PP1), iii) Wiscott-Aldrich homology 2 (WH2) domain and iv) β-sheets within the CARP domain. Further, we deleted the last four aa residues (Δ4CT), which have been implicated in actin dynamics recently (Kotila et al., 2018). Similar to WT-CAP1 and both deletion constructs, all mutant CAP1 variants were located in growth cones (Fig. S3). Expression of CAP1-PP1, CAP1-WH2, CAP1-β-sheet or CAP1-Δ4CT normalized growth cone size in CAP1-KO neurons (PP1: 16.96±2.04, n=50/3, P<0.001; WH2: 21.88±1.85, n=50/3, P<0.001; CAP1-β-sheet: 29.40±2.43, n=75/3, P<0.001; CAP1-Δ4CT: 20.38±1.44, n=80/3, P<0.001). Instead, CAP1-HFD expression only slightly reduced growth cone size in CAP1-KO neurons (34.73±1.97, n=100/3, P<0.05), which was still larger when compared to CTR neurons or WT-CAP1-expressing CAP1-KO neurons (P<0.001 and <0.05, respectively). Together, our data identified HFD to be relevant for CAP1 in growth cones. Since previous studies revealed an interaction of HFD with ADF/cofilin-actin-complexes (Kotila et al., 2019; Shekhar et al., 2019), we hypothesized that CAP1 interacts with ADF/cofilin in growth cones.

**Figure 6.**
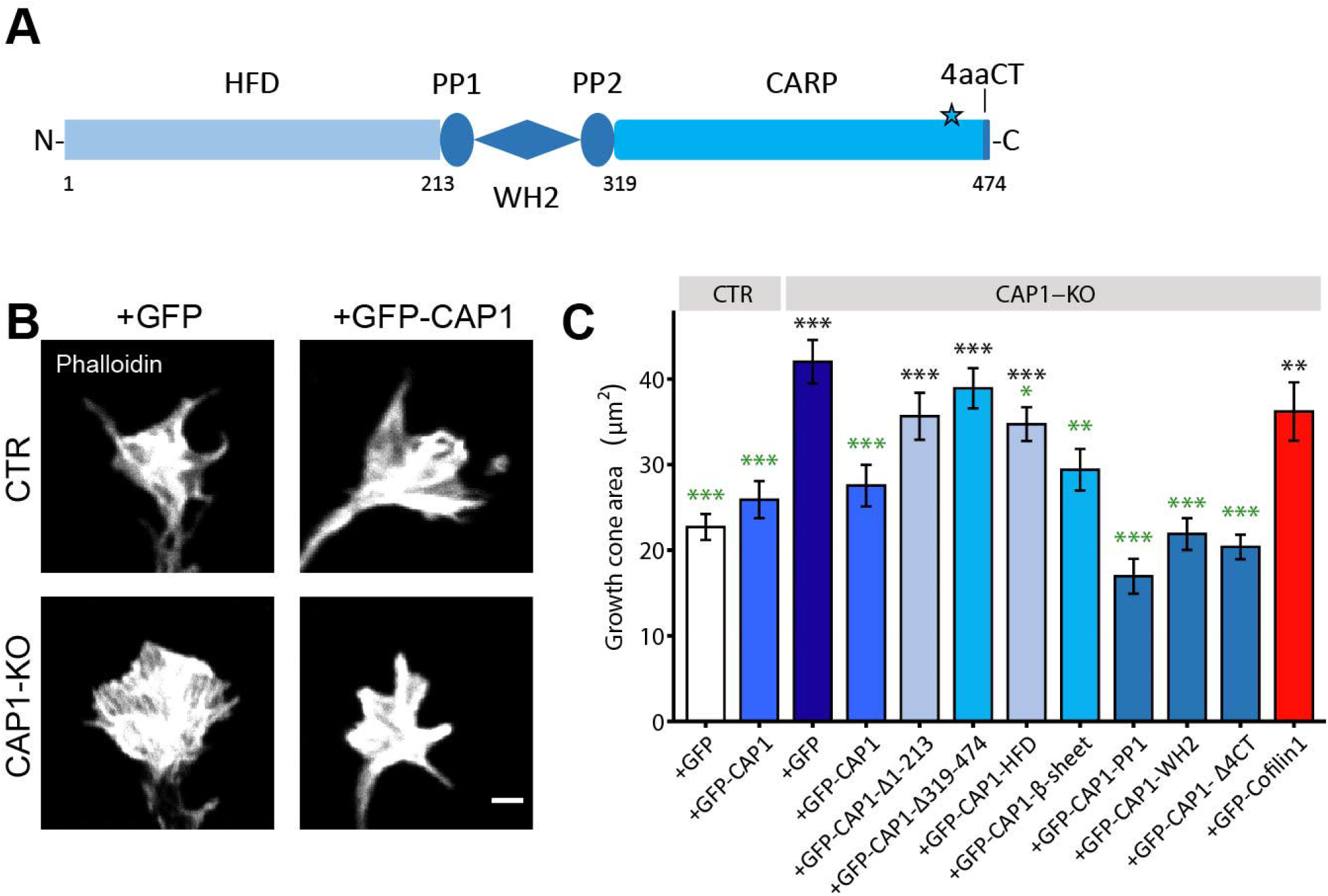
Helical-folded domain is relevant for CAP1 function in growth cones. **(A)** Scheme showing protein domains of CAP1. HFD: helical folded domain, PP1: proline-rich stretch 1, WH2: Wiscott-Aldrich homology 2 domain, PP2: proline-rich stretch 2, CARP domain, 4aaCT: last four C-terminal amino acid residues. Asterisk marks β-sheet mutation within CARP domain. **(B)** Representative micrographs of phalloidin-stained growth cones from stage 2 CTR and CAP1-KO neurons expressing GFP or GFP-CAP1. **(C)** Growth cone size in CTR and CAP1-KO neurons that express either GFP, WT-CAP1, CAP1 deletion constructs (CAP1-Δ1-213, CAP1-Δ319-474), CAP1 mutant constructs or WT-cofilin1. Δ4CT: CAP1 deletion mutant lacking C-terminal four amino acid residues. Black asterisks indicate conditions significantly different from GFP-expressing CTR neurons, green asterisks indicate conditions significantly different from GFP-expressing CAP1-KO neurons. Scale bars (in µm): 2 (B); **: P<0.01, ***: P<0.001.

### Acute CAP1 inactivation impaired growth cone size and actin dynamics

The ADF/cofilin family comprises the family members ADF, cofilin1 and cofilin2. All are expressed in the mouse brain, but analyses of gene targeted mice identified cofilin1 as the major family member during brain development (Bellenchi et al., 2007; Görlich et al., 2011; Flynn et al., 2012; Gurniak et al., 2014). Moreover, cofilin1 has been located in growth cones and implicated in actin dynamics downstream of guidance cues (Dent et al., 2011; Flynn et al., 2012; Garvalov et al., 2007; Wang et al., 2016). We therefore hypothesized that CAP1 cooperates with cofilin1 in growth cones. To test this hypothesis, we chose a genetic approach and compared under identical experimental conditions growth cones from single mutant neurons lacking either CAP1 or cofilin1 or both ABP. To do so, we isolated hippocampal neurons from i) CAP1^flx/flx^ mice, ii) Cfl1^flx/flx^ mice (Bellenchi et al., 2007) and iii) double transgenic mice (CAP1^flx/flx^/Cfl1^flx/flx^) at E18.5. Gene inactivation in isolated neurons was achieved by expression of mCherry-tagged Cre (mC-Cre), neurons expressing a catalytic inactive mCherry-Cre (mC-Cre-mut) served as controls (Kullmann et al., 2020b). Immunocytochemistry against CAP1 confirmed inactivation upon mC-Cre expression, but not upon mC-Cre-mut expression (Fig. S5). We reset neurons into an undifferentiated stage by re-plating them at DIV2 (Biswas and Kalil, 2018), and investigated growth cones 24 h later. First, we tested whether acute CAP1 inactivation compromised actin dynamics in growth cones. To do so, we performed FRAP experiments in GFP-actin-expressing neurons, and we determined retrograde F-actin flow in LifeAct-GFP-expressing neurons, similar to our analyses in CAP1-KO neurons (Fig. 4). Compared to mC-Cre-mut-expressing control neurons, actin turnover rate (expressed as third-recovery time (t_1/3_)) was reduced and stable actin fraction was increased in mC-Cre-expressing CAP1^flx/flx^ neurons (Fig. 7A-D, Movies S7-8; t_1/3_ (in s): mC-Cre-mut: 27.89±2.87, mC-Cre: 47.75±8.20, n=19/3, P<0.05; stable fraction: mC-Cre-mut: 0.74±0.03, mC-Cre: 0.56±0.04, n=19/3, P<0.001).

**Figure 7.**
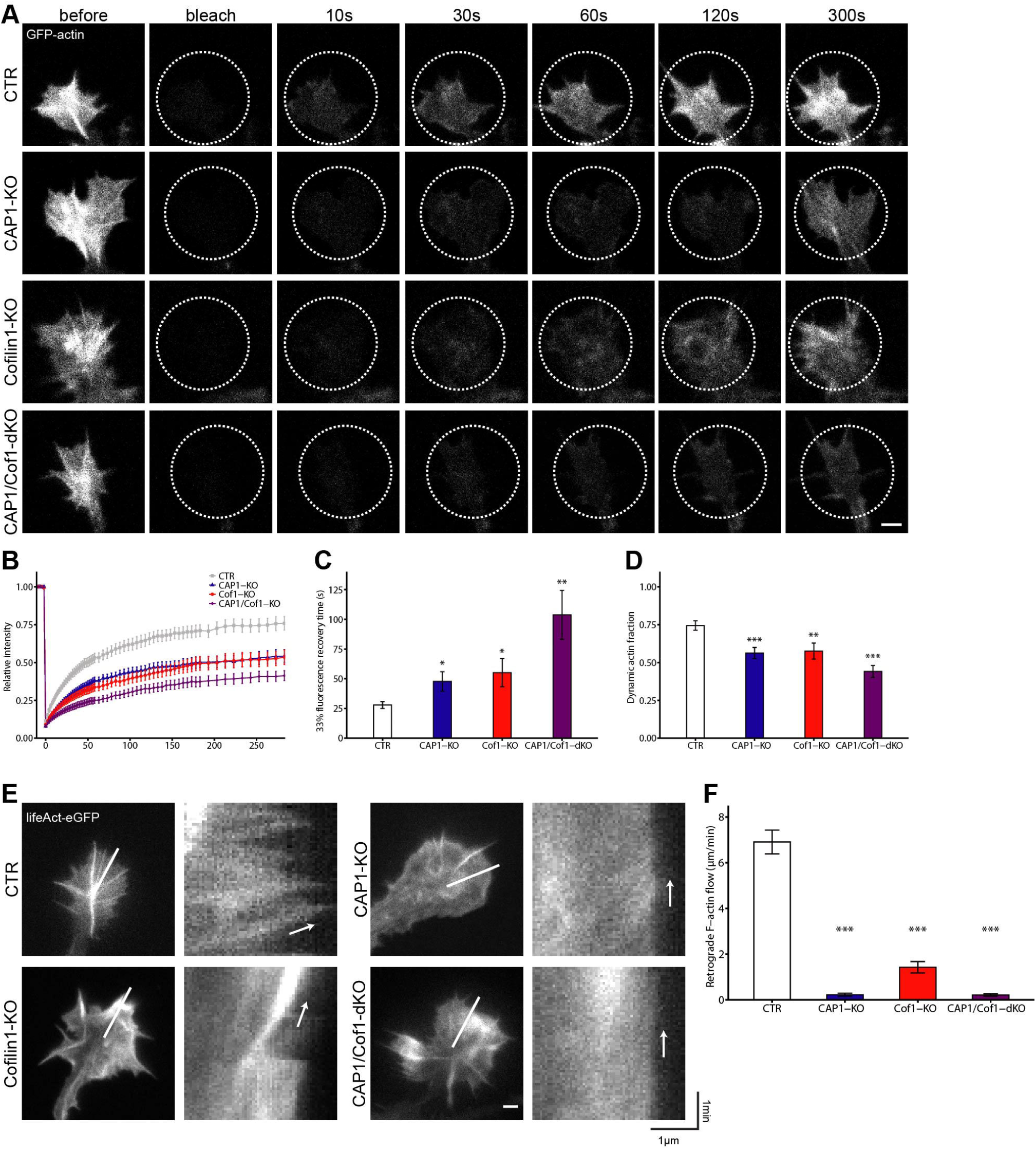
Impaired actin dynamics in neurons lacking either CAP1, cofilin1 or both ABPs. **(A)** Image sequence of GFP-actin in growth cones from control neurons expressing catalytic inactive Cre (mC-Cre-mut) or neurons from CAP1^flx/flx^, Cfl1^flx/flx^, and CAP1^flx/flx^/Cfl1^flx/flx^ mice expressing catalytic active Cre (mC-Cre). Dashed lines encircle bleached areas. **(B)** Recovery curves for GFP-actin in growth cones from control and mutant neurons. **(C)** Third-recovery time for GFP-actin in growth cones from control and mutant neurons. **(D)** Dynamic actin fraction in growth cones from control and mutant neurons. **(E)** Representative micrographs of growth cones from LifeAct-GFP-expressing control and mutant neurons. Lines indicate where kymographs (shown on the right) have been generated from. Arrows indicate the retrograde F-actin flow. **(F)** Velocity of retrograde F-actin flow in growth cones from control and mutant neurons. Scale bars (in µm): 2 (A, E); *: P<0.05, **: P<0.01, ***: P<0.001.

Likewise, retrograde F-actin flow was reduced in mC-Cre-expressing CAP1^flx/flx^ neurons (Fig. 7E-F, Movies S9-10; (in µm/min) mC-Cre-mut: 6.91±0.52, mC-Cre: 0.22±0.07, n=32/3, P<0.001). Impaired actin dynamics in growth cones from mC-Cre-expressing CAP1^flx/flx^ neurons was associated with roughly 40% increased growth cone size (Fig. 8A-B; (in µm^2^) mC-Cre-mut: 20.07±0.73, mC-Cre: 27.72±2.12, n=90/3, P<0.05). Together, acute CAP1 inactivation in hippocampal neurons caused growth cone defects similar to those we reported for CAP1-KO neurons, thereby proving suitability of the experimental approach.

**Figure 8.**
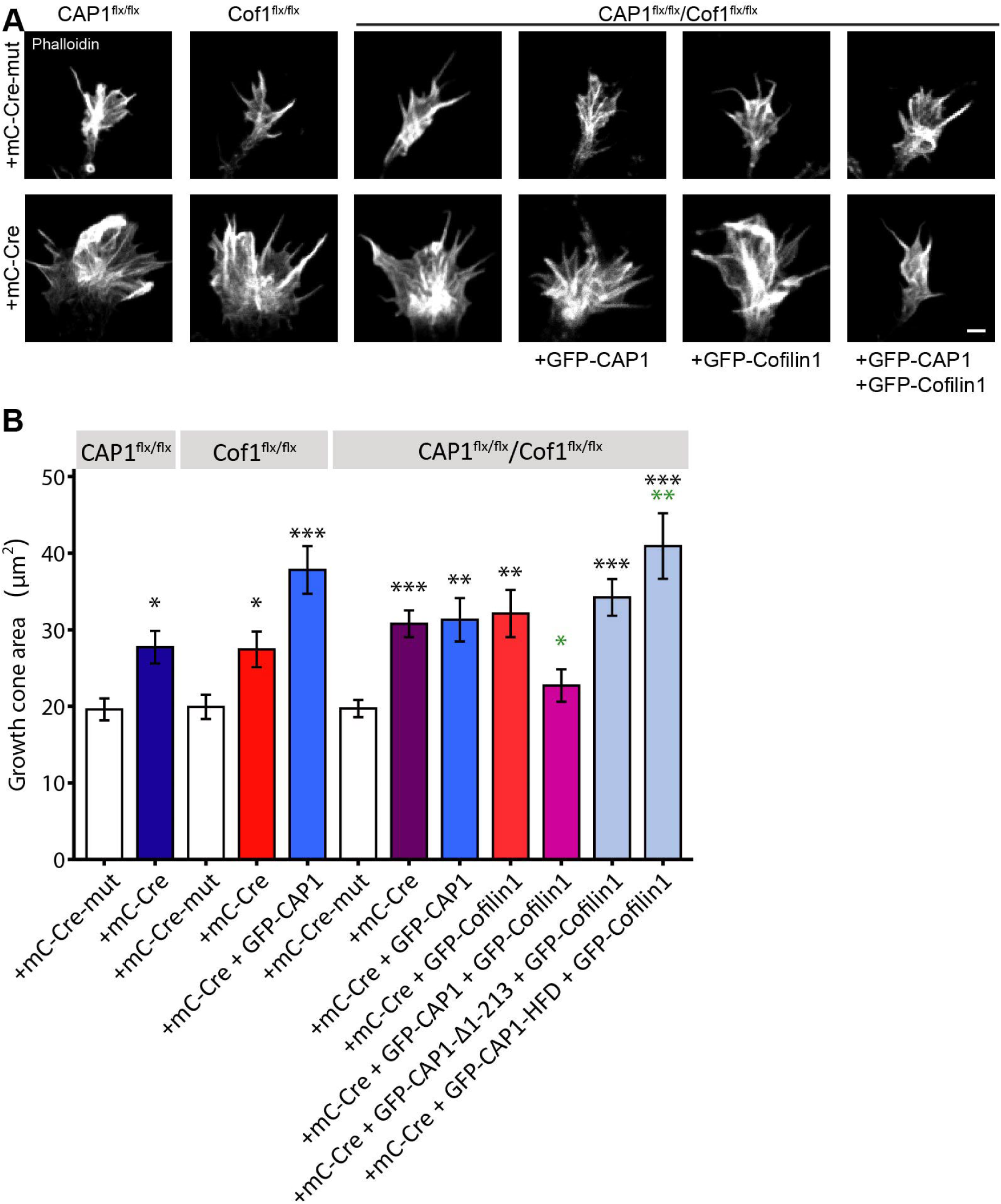
CAP1 and cofilin1 synergistically control growth cone size. **(A)** Representative micrographs of phalloidin-stained growth cones from CAP1^flx/flx^ and Cfl1^flx/flx^ neurons expressing either catalytic inactive Cre (mC-Cre-mut) or catalytic active Cre (mC-Cre) as well as from CAP1^flx/flx^/Cfl1^flx/flx^ neurons expressing mC-Cre-mut or mC-Cre alone or together with GFP-CAP1, GFP-Cofilin1 or both ABPs. **(B)** Growth cone size in CAP1^flx/flx^, Cfl1^flx/flx^, or CAP1^flx/flx^/Cfl1^flx/flx^ neurons expressing either mC-Cre-mut or mC-Cre as indicated, and in mC-Cre-expressing Cfl1^flx/flx^ and CAP1^flx/flx^/Cfl1^flx/flx^ neurons that additionally express the indicated constructs. Black asterisks indicate conditions significantly different from GFP-expressing CTR neurons, green asterisks indicate conditions significantly different from GFP-expressing CAP1-KO neurons. Scale bars (in µm): 2 (A); n.s.: P≥0.05, *: P<0.05, **: P<0.01, ***: P<0.001.

### Cofilin1 inactivation caused growth cone defects similar to CAP1 inactivation

Next, we performed identical experiments with neurons from Cfl1^flx/flx^ mice. Compared to mC-Cre-mut-expressing control neurons, actin dynamics in growth cones was impaired upon mC-Cre expression in Cfl1^flx/flx^ neurons as judged from reduced actin turnover and increased stable actin fraction in FRAP experiments (Fig. 7A-D, Movie S11; t_1/3_ (in s): 55.17±11.94, n=21/3, P<0.05; stable fraction: 0.58±0.05, n=21/3, P<0.01) as well as from reduced retrograde F-actin flow (Fig. 7E-F, Movie S12; (in µm/min) 1.42±0.25, n=30/3, P<0.001). Hence, actin dynamics in growth cones was similarly impaired upon acute inactivation of either CAP1 or cofilin1. Moreover, growth cones were enlarged by roughly 40% upon cofilin1 inactivation, similar to CAP1 inactivation (Fig. 8A-B; (in µm^2^) 27.43±2.32, n=70/3, P<0.05). Interestingly, overexpression of WT-CAP1 did not reduced growth cone size in cofilin1 mutants ((in µm^2^) 37.81±3.11, n=55/3, P<0.001), similar to WT-cofilin1 overexpression in CAP1-KO neurons (Fig. 6C; (in µm^2^) 36.20±3.41, n=50/3, P=0.23). Hence, acute inactivation of CAP1 and cofilin1 induced similar defects in growth cone actin dynamics and size, demonstrating that they are of equal importance in growth cones. These findings support our hypothesis that both ABPs act together in growth cones.

### CAP1 cooperates with cofilin1 in growth cones

To ultimately test this hypothesis, we examined growth cones in double mutant neurons (mC-Cre-expressing CAP1^flx/flx^/Cfl1^flx/flx^ neurons). FRAP experiments revealed strongly reduced actin turnover in double mutant neurons when compared to mC-Cre-mut-expressing control neurons (Fig. 7A-C, Movie S13; t_1/3_ (in s) 103.77±20.56, n=22/3, P<0.01). Reduced actin turnover was associated with an increased fraction of stable actin (Fig. 7D; 0.44±0.04, n=22/3, P<0.001). Likewise, retrograde F-actin flow was strongly reduced in double mutant neurons (Fig. 7E-F; Movie S14; (in µm/min) 0.21±0.06, n=30/3, P<0.001). Further, we found growth cone size increased by roughly 50% in double mutant neurons when compared mC-Cre-mut-expressing CAP1^flx/flx^/Cfl1^flx/flx^ controls (Fig. 8A-B; (in µm^2^) 30.78±1.74, n=80/3, P<0.001). Growth cone size in double mutant neurons was not different from CAP1 or cofilin1 single mutant neurons (P=0.27 or 0.27, respectively). Notably, growth cone size in double mutant neurons was unchanged upon expression of either WT-CAP1 or WT-cofilin1 ((in µm^2^) WT-CAP1: 31.30±2.84, n=70/3, P=0.89; WT-cofilin1: 32.12±3.08, n=100/3, P=0.69). Instead, compound expression of WT-CAP1 and WT-cofilin1 normalized growth cone size in double mutant neurons ((in µm^2^) 22.71±2.13, n=90/3, P<0.05), and growth cone size in these neurons was not different from mC-Cre-mut-expressing control neurons (P=0.39). While expression of WT-cofilin1 together with WT-CAP1 normalized growth cone size in double mutant neurons, expression of WT-cofilin1 together with CAP1 variants either possessing a mutated HFD (CAP1-HFD) or lacking the HFD (CAP1-Δ1-213) failed in rescuing growth cone size ((in µm^2^) CAP1-HFD: 40.93±4.29, n=80/3, P<0.01; CAP1-Δ1-213: 34.22±2.40, n=120/3, P=0.28). Together, these data demonstrated that CAP1 and cofilin1 cooperate in growth cones and that CAP1’s HFD is crucial for this interaction.

## Discussion

The present study aimed at deciphering the role of CAP1 for mammalian brain development, which we found expressed during neuron differentiation and enriched in growth cones. By generating a conditional mouse strain and by exploiting primary hippocampal neurons from brain specific mutant mice, we identified CAP1 as a novel regulator of F-actin organization and dynamics in growth cones, which was relevant for growth cone morphology, motility and response to guidance cues. Growth cone defects in CAP1 mutant neurons were associated with retarded neuron differentiation and impaired neuron connectivity in CAP1 mutant brains. Rescue experiments in CAP1 mutant neurons, and the analysis of double mutant neurons lacking CAP1 and cofilin1 revealed a cooperation of both ABP in neuronal actin dynamics and growth cone function.

While CAPs have been recognized as ABPs almost two decades ago (Balcer et al., 2003; Bertling et al., 2004; Freeman and Field, 2000; Hubberstey and Mottillo, 2002), significant progress in their molecular functions has been achieved just recently (Jansen et al., 2014; Johnston et al., 2015; Kotila et al., 2018; Kotila et al., 2019; Mu et al., 2019; Shekhar et al., 2019). These studies, which either exploited recombinant proteins or have been performed in yeast or non-neuronal cell lines, implicated CAP1 and CAP2, but also their yeast homolog Srv2 (suppressor of Ras2-Val19) in actin dynamics. In good agreement with these studies, our live cell imaging data unraveled a function for CAP1 in actin dynamics of growth cones. However, despite the substantial advancements of their molecular activities, the physiological functions of mammalian CAPs largely remained elusive, also because appropriate mouse models were lacking. This holds true specifically for CAP1 (Jang et al., 2019), while recent mouse studies implicated CAP2 in heart physiology, myofibril differentiation and dendritic spine morphology (Peche et al., 2012; Field et al., 2015; Kepser et al., 2019; Pelucchi et al., 2020). Our conditional mouse model allowed us to study the brain function of CAP1. We found that CAP1 controls actin dynamics in differentiating neurons, which was relevant for growth cone morphology and function. We showed that growth cone motility, and consequently its exploratory behavior, was severely disturbed upon CAP1 inactivation and that CAP1 mutant growth cones were strongly impaired in responding to attractant and repellent cues. As expected for defective growth cone actin dynamics (Gomez and Letourneau, 2014), neurite formation and morphology was altered in CAP1 mutant neurons, and neuron connectivity was compromised in CAP1 mutant brains. Instead, neuron layering and brain anatomy was largely preserved in CAP1 mutant mice, demonstrating that CAP1 was dispensable for various important aspects of brain development such as neural stem cell proliferation and differentiation, neuron production and migration or layer formation in the cerebral cortex – different from other ABP that control neuronal actin dynamics during brain development (Bellenchi et al., 2007; Flynn et al., 2012; Kullmann et al., 2020a). Hence, our data led us suggest that CAP1 acquired a specific function in neuron differentiation and in establishing neuron connectivity. The latter is in good agreement with a previous study that implicated the Drosophila homolog capulet (also known as act up) in axonal midline crossing (Wills et al., 2002).

Although numerous ABPs have been implicated in growth cone motility (Dent et al., 2011), knowledge about the mechanisms that control actin dynamics in growth cones is still fragmented (Gomez and Letourneau, 2014; Omotade et al., 2017). Our data add CAP1 to the list of actin regulators relevant for growth cone function. By live cell imaging, we showed that CAP1 is relevant for dynamizing the growth cone actin cytoskeleton, and our pharmacological data revealed that CAP1 acts downstream of guidance cues such as BDNF, EphA5, Sema4D or Slit-1. Further, our rescue experiments identified CAP1’s HFD to be relevant in growth cones. Instead, expression of mutant CAP1 variants i) in which either the PP1, the WH2 domain or a β-sheet within the CARP domain were mutated or ii) which lacked the C-terminal four aa residues restored growth cone defects in CAP1 mutant neurons, very similar to WT-CAP1. We therefore concluded that CAP1’s interaction with ADF/cofilin-actin complexes is crucial in growth cones (Kotila et al., 2019; Moriyama and Yahara, 2002; Quintero-Monzon et al., 2009; Shekhar et al., 2019). Instead, CAP1’s nucleotide exchange activity on G-actin that depends on its CARP and WH2 domains as well as C-terminal four aa residues seems to be dispensable in growth cones, very much alike CAP1’s interaction with actin regulators such as profilin or Abl1 that depends on PP1 (Bertling et al., 2007; Chaudhry et al., 2010; Kotila et al., 2018; Makkonen et al., 2013; Mattila et al., 2004; Nomura and Ono, 2013). Indeed, we confirmed the relevance of the CAP1-cofilin1 interaction in growth cones in a genetic approach, in which i) we found similar defects in growth cone actin dynamics upon inactivation of either CAP1, cofilin1 or both ABPs, ii) we found a similar increase in growth cone size in both single mutant neurons and double mutant neurons, iii) growth cone size in double mutant neurons was normalized upon expression of CAP1 together with cofilin1, but not upon expression of CAP1 or cofilin1 alone, and iv) growth cone size in double mutant neurons was not normalized upon expression of cofilin1 together with CAP1 variants possessing a mutated HFD. We therefore conclude that i) CAP1 cooperates with cofilin1 in growth cone actin dynamics and ii) CAP1 is mandatory for cofilin1 activity in growth cones and *vice versa*. Cofilin1 is a key regulator of the neuronal actin cytoskeleton (Rust, 2015). By exploiting gene targeted mice, we and others previously demonstrated that cofilin1 is important for brain development and function (Bellenchi et al., 2007; Flynn et al., 2012; Rust et al., 2010; Wolf et al., 2015; Zimmermann et al., 2015). Moreover, several studies implicated cofilin1 in growth cone motility and revealed a complex and context-dependent function in growth cone dynamics (Dent et al., 2011; Gomez and Letourneau, 2014; Omotade et al., 2017; Wang et al., 2016; Zhang et al., 2019). Very similar to CAP1, cofilin1 acts downstream of repellent cues in growth cone collapse (Grintsevich et al., 2016; Hsieh et al., 2006; Piper et al., 2006; Wen et al., 2007). Hence, a cooperation of CAP1 and cofilin1 as demonstrated here is in line with these studies. Moreover, it fits well to previous studies that either exploited recombinant proteins or have been performed in yeast or non-neuronal cell lines (Bertling et al., 2004; Jansen et al., 2014; Johnston et al., 2015; Kotila et al., 2019; Moriyama and Yahara, 2002; Quintero-Monzon et al., 2009; Shekhar et al., 2019). These studies revealed that CAP1 and cofilin1 synergistically control actin dynamics and that CAP1 releases cofilin1 from its complex with G-actin, thereby allowing a new cycle of actin depolymerization. However, these study further implicated CAP1 in nucleotide exchange on G-actin that recycles G-actin for F-actin assembly. Our data suggest that G-actin recycling in growth cones does not depend on CAP1 and might be carried out by other ABPs such as profilin, for which such a function in growth cones has been proposed earlier (Gomez and Letourneau, 2014; Omotade et al., 2017; Vitriol and Zheng, 2012).

In summary, we identified CAP1 as a novel regulator of actin dynamics in growth cones that is relevant for growth cone function, neuron differentiation and neuron connectivity in the mouse brain and that cooperates with cofilin1 in neuronal actin dynamics and growth cone function.

## Material and Methods

### Transgenic mice

Brain-specific deletion of CAP1 was achieved by crossing mice carrying conditional CAP1 alleles (CAP1^flx/flx^) with CAP1^+/flx^ mice that additionally express a Cre transgene under control of the nestin promoter (Tronche et al., 1999). Cfl1^flx/flx^ mice have been generated previously (Bellenchi et al., 2007) and CAP1^flx/flx^/Cfl1^flx/flx^ mice were generated by crossing a CAP1^flx/flx^ mice with Cfl1^flx/flx^ mice. Mice were housed in the animal facility of the University of Marburg on 12-hour dark-light cycles with food and water available *ad libitum*. Treatment of mice was in accordance with the German law for conducting animal experiments and followed the guidelines for the care and use of laboratory animals of the U.S. National Institutes of Health. Killing of mice has been approved by internal animal welfare authorities at Marburg University (references: AK-5-2014-Rust, AK-6-2014-Rust, AK-5-2018-Rust). Generation and breeding of gene targeted mice has been approved by the Regierungspräsidium Giessen (references: V54-19c2015h01MR20/30 #83/2015, V54-19c2015h01MR20/29 #G22/2016) and by the Thüringer Landesamt für Verbraucherschutz (222684-04-02-060/14).

### Cell culture and transfection

Primary hippocampal neurons from embryonic day 18 (E18) mice were prepared as previously described (Antoniou et al., 2018). Briefly, hippocampi were dissociated individually to keep their genetic identity and seed with a density of 31,000 neurons per cm^2^ on 0.1 mg/ml poly-L-lysine-coated coverslips (15,500 neurons for each genotype per cm^2^ in mixed cultures). Neurons were maintained for 5 h to 3 d in a humidified incubator at 37°C with 5% CO_2_ in Neurobasal medium containing 2% B27, 2 mM GlutaMax, 100 µg/ml streptomycin, and 100 U/ml penicillin (Gibco, Invitrogen). For overexpression of plasmids, neurons were nucleofected prior to plating with Amaxa nucleofector system (Lonza) according to manufacturer’s protocol. For each nucleofection, 3 µg plasmid was transfected into 250,000 neurons, which were then plated at a density of 66,000 neurons per cm^2^. For re-plating, neurons were seeded in wells coated with 0.05 mg/ml poly-L-lysine and detached with TrypLE™ Express (Gibco) at DIV2, similar to previous studies (Biswas and Kalil, 2018). Thereafter, neurons were pelleted at 7,000 rpm for 7 min, plated on 0.1 mg/ml poly-L-lysine-coated coverslips and fixed 24 h later. Following constructs were used: GFP-CAP1 and mutant GFP-CAP1 and deletion myc-CAP1 variants (GenScript), GFP (GenScript), GFP-actin and LifeAct-GFP (Robert Grosse lab), GFP-cofilin1 (Rehklau et al., 2017), mCherry-Cre and mCherry-Cre-mutant (Kullmann et al., 2020b).

### Immunocytochemistry

Cultured neurons were fixed for 10 min in 4% paraformaldehyde (PFA) in PBS under cytoskeleton preserving conditions. After 5 min incubation with 0.4% gelatin with 0.5% Triton-X100 in PBS (carrier solution), neurons were incubated with following primary antibodies (in carrier solution): mouse anti-CAP1 (1:100, Abnova), rabbit anti-doublecortin (1:500, Abcam), rabbit anti-GFP (1:1,000, ThermoFisher Scientific), mouse anti tau-1 (1:200, Merck Millipore), mouse anti-c-Myc (1:200, ThermoFisher Scientific) and chicken anti-mCherry (1:500, Abcam). Thereafter, neurons were washed in PBS and incubated with the following secondary antibodies (in carrier solution): anti-mouse and anti-rabbit IgG coupled to either AlexaFluor488, AlexaFluor546 or AlexaFluor647 (1:500, Invitrogen) and AlexaFluor555-coupled anti-chicken IgG (1:500, Invitrogen). F-actin was visualized by staining with phalloidin coupled to either AlexaFluor488 (1:100, Invitrogen) or AlexaFluor647 (1:100, Cell Signaling Technologies). In each experiment, neurons were stained with the DNA dye Hoechst (1:1,000, Invitrogen). Image acquisition was done with a Leica TCS SP5 II confocal microscope setup, and image analysis was performed with ImageJ (Schindelin et al., 2012).

### Growth cone morphology

Growth cone morphology was assessed by determining growth cone solidity (growth cone area divided by hull area), similar to previous studies (Chitsaz et al., 2015). ‘Hand-shaped’ growth cones were defined as growth cones with visible filopodia protruding out of the lamellipodium, while smooth growth cones were lacking these protruding filopodia.

### Live cell imaging

For fluorescence recovery after photobleaching (FRAP), GFP-actin transfected neurons were seeded in a 22 mm glass-bottom dish either directly after nucleofection or after re-plating and cultured for 1 d. Neurons were imaged with a Leica TCS SP5 II in a chamber heated to 35°C and the medium was exchanged with CO_2_-saturated HBS solution. For pre-bleaching condition, 5 images were acquired and in total 65 images for the fluorescence recovery over a time course of 5 min. Images were analyzed with ImageJ (Schindelin et al., 2012) and recovery curve calculated with R. The retrograde F-actin flow was measured on neurons, which were transfected with LifeAct-GFP and cultured in a 22 mm glass-bottom dish either directly after nucleofection or after re-plating for 1 d. Neurons were imaged in CO_2_-regulated incubation chamber maintained at 37°C with Leica DMi8 Thunder microscope system and a Leica DFC9000 GTC camera. Images were acquired every 5 s for 5 min. Generation and analysis of kymograph was performed with ImageJ (Schindelin et al., 2012). For DIC imaging, neurons were cultured as described above and imaged in a chamber maintained at 37°C and the medium was exchanged with CO_2_-saturated HBS solution. Images were acquired every 5 s for 10 min at a Leica DMi8 setup.

### Growth cone collapse assay

Neurons were cultured for 1 d before incubation with 100 µM BDNF (PeproTech), 1 µg/µl Ephrin A5 (R&D System), 50 nM Semaphorin 4D (Thomas Worzfeld lab) or 1 µg/µl Slit-1 (R&D System) for 60 min. Neurons were imaged with a Leica TCS SP5 II microscope system and analyzed with ImageJ (Schindelin et al., 2012). Collapsed and non-collapsed growth cones were defined according to previous studies (Muller et al., 1990). For BDNF-treated neurons only growth cone size was measured.

### dSTORM

Neurons were plated in a µ-Slide 8 Well chamber (ibidi) at a density of 30,000 cells per well. 24 h after plating, neurons were fixed with 1% PFA and 0.05% Triton-X for 1 min followed by 3% PFA for 10 min. After washing, samples were quenched with 1 mg/ml NaBH_4_ for 10 min and washed again. For staining, cells were blocked with 100 µM L-lysine in ImageiT™ FX Signal Enhancer solution (Thermo Fisher Scientific) for 1 h and then stained with Phalloidin-AF647 (1:50 in PBS, Cell Signal Technology) for 48 h at 4 °C. Neurons were post-fixed with 4% PFA for 10 min, washed with 0.05% Tween20 in PBS and incubated again with Phalloidin-AF647 (1:50 in PBS) for 24 h at 4 °C. Neurons were post-fixed for a second time with 4% PFA for 10 min, washed with 0.05% Tween20 in PBS and stored in PBS at 4 °C until imaging.

Image acquisition was carried out on a customized single-molecule localization microscope as described before (Virant et al., 2018). Before imaging, samples were incubated with infrared beads (FluoSphere infrared fluorescent Carboxylate-Modified Microspheres, ThermoFisher, USA; ex/em 715/755□nm; 1:2,000 in PBS) for 10 min and then PBS was exchanged with dSTORM buffer containing 100 mM mercaptoethylamine (MEA) with a glucose oxygen scavenger system (van de Linde et al., 2011). The sample was illuminated in HILO (Highly Inclined and Laminated Optical sheet) mode and growth cones were recorded at 20Hz for 30,000 frames with a final intensity of the 640 nm laser adjusted to 1-2 kW cm^-2^ adapted from the parameters of the dSTORM recordings (Baarlink et al., 2017; Tokunaga et al., 2008). Recorded fluorescent single molecule spots from the image acquisition were localized using ThunderSTORM (Ovesny et al., 2014) and further processed using customized Python scripts, kindly provided by Dr. Bartosz Turkowyd. The localization files were filtered for out-of-focus signals (80 nm < PSF sigma < 200 nm, uncertainty in xy < 50 nm) and corrected for sample drift during image acquisition using the infrared beads in the sample as described before (Balinovic et al., 2019). The final experimental localization precision in the processed localization files was determined by calculating the NeNA precision value (Endesfelder et al., 2014). Using RapidSTORM 3.0 (Wolter et al., 2012), super-resolved images were reconstructed from the localization files with a pixel size of 10 nm and were overlaid with a Gaussian blur according to the individual NeNA localization precision using ImageJ (Schindelin et al., 2012). The number of phalloidin-AlexaFluor647 localizations per growth cone area was measured with the software swift (written in C++, Endesfelder group, unpublished) by selecting the growth cone and normalized to the growth cone area. Filopodia were counted with ImageJ (Schindelin et al., 2012).

### Protein analysis

Cortices of E18.5 mice were homogenized in 500 µl RIPA buffer containing 50 mM Tris HCl (pH7.5), 150 mM NaCl, 0.5% NP40, 0.1% SDS and protease inhibitor (Complete, Roche). After centrifugation at 14,000 rpm for 10 min at 4°C, samples were boiled for 5 min at 95°C in Laemmli buffer including 6% DTT. Equal protein amounts were separated by SDS PAGE and blotted onto a polyvinylidene difluoride membrane (Merck) by using a Wet/Tank Blotting System (Biorad). Membranes were blocked in Tris-buffered saline (TBS) containing 5% milk powder, 0.5% Tween-20 and 0.02% sodium acid for 1 hour and afterwards incubated with primary antibodies in blocking solution over night at 4°C. As secondary antibodies, horseradish peroxidase (HRP)-conjugated antibodies (1:20,000, Thermo Fisher Scientific) were used and detected by chemiluminescence with ECL Plus Western Blot Detection System (GE Healthcare). Following primary antibodies were used: mouse anti-CAP1 (1:1,000, Abnova), mouse anti-GAPDH (1:1,000, R&D System). For secondary antibody, anti-mouse and anti-rabbit coupled to HRP (Thermo Fisher Scientific) were used and the blots were developed with ECL Prime Western Blotting Detection Reagent (GE Healthcare).

### Histology and immunohistochemistry

Nissl staining and immunohistochemical staining was performed as described previously (Kullmann et al., 2020a). Briefly, E18.5 mice were killed by decapitation and brains were fixed for 2 h in PBS containing 4 % PFA. Thereafter, 25 µm transversal brain sections were generated by a Leica CM3050 S cryostat. For immunohistochemistry, brain sections were incubated for 1 h with 2% BSA, 3% goat serum, 10% donkey serum and 0.5% NP40 in PBS and stained over night at 4°C with following primary antibodies in 2% BSA and 0.5% NP40 in PBS: rat anti-Ctip2 (1:1,000, Abcam), rabbit anti-Neurofilament200 (1:80, Sigma-Aldrich), rabbit anti-Tbr1 (1:200, Abcam). As secondary antibodies, AlexaFluor488 anti-rabbit, AlexaFluor488 anti-rat and AlexaFluor546 anti-rabbit IgG was used. For Nissl staining, brains sections were incubated in staining solution according to the manufacturer’s instructions. Image acquisition was done with a Leica TCS SP5 II confocal microscope setup and Leica M80 equipped with a Leica DFC295 camera.

## Statistical Analysis

Statistical tests were done in R or Sigma Plot. For comparing mean values between groups, student’s t-test or Mann-Whitney U-test was performed. Analyzing the rescue conditions, ANOVA with post-hoc test was used. Stage distribution (‘hand shaped’ vs. smooth, non-collapsed vs. collapsed) was tested for differences with χ2-test.

## Acknowledgements

We thank Renate Gondrum and Katrin Schorr for excellent technical support and Ralf Jacob and the Bioimaging Core Facility of the University of Marburg for support in live imaging. This work was supported by research grants from the Deutsche Forschungsgemeinschaft (DFG) to MBR (RU 1232/7-1). FS received a fellowship from the University of Marburg and was funded by the DFG Research Training Group ‘Membrane Plasticity in Tissue Development and Remodeling’ (GRK 2213). We thank Robert Grosse (University of Freiburg) for GFP-actin and LifeAct-GFP constructs, Thomas Worzfeld (University of Marburg) for recombinant Semaphorin 4D, David Solecki (St. Jude Children’s Research Hospital, Memphis) for mCherry-Cre constructs and Walter Witke (University of Bonn) for GFP-cofilin1 constructs and Cfl1^flx/flx^ mice.

## Author contributions

Blastocyst injection of CAP1 targeted ES cells was performed by CAH. Experiments were designed and results were discussed by FS, TAD, IM, JW, UE and MBR. Data were analyzed by FS, TAD, IM, JW, UE and MBR. Manuscript was written by MBR and FS.

## Legends to supplementary figures

**Figure S1.**
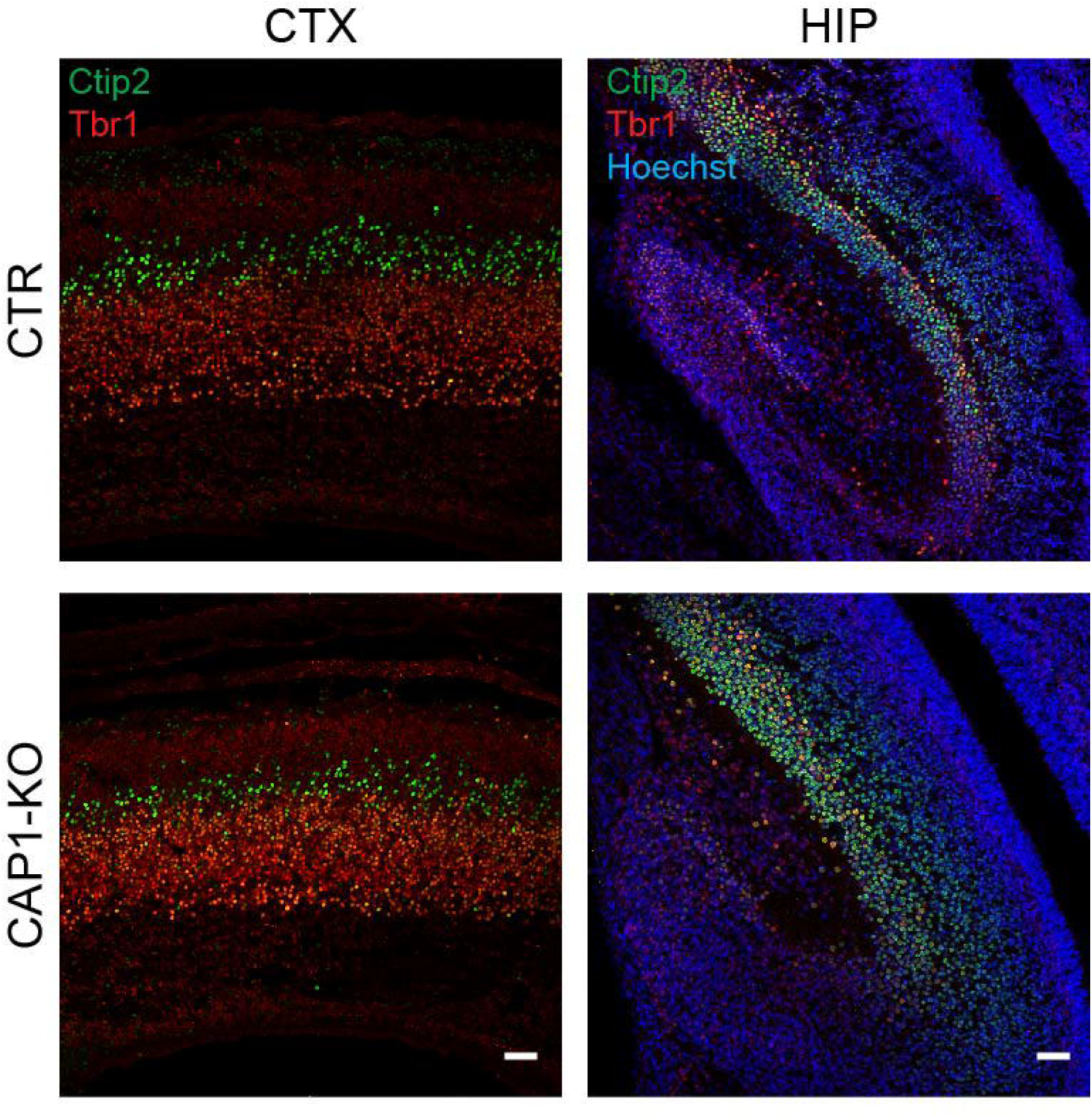
Antibody staining of transversal brain sections from E18.5 CTR and CAP1-KO mice against the neuronal markers Tbr1 (red) and Ctip2 (green). CTX: cerebral cortex, HIP: hippocampus. Sections were counterstained with the DNA dye Hoechst (blue). Scale bars (in µm): 50.

**Figure S2.**
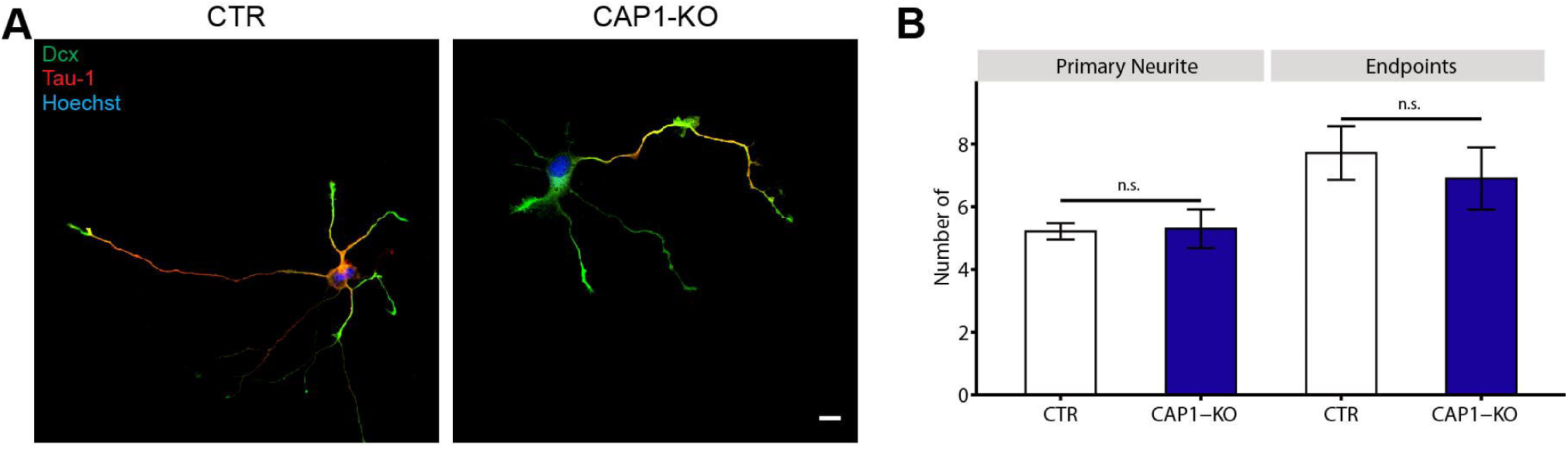
**(A)** Stage 3 neurons stained with antibodies against the axon marker tau-1 (red) and Dcx (green). Neurons were counterstained with Hoechst (blue). **(B)** Number of primary neurites and neurite endpoints in stage 3 CTR and CAP1-KO neurons. Scale bar (in µm): XY. n.s.: P≥0.05.

**Figure S3.**
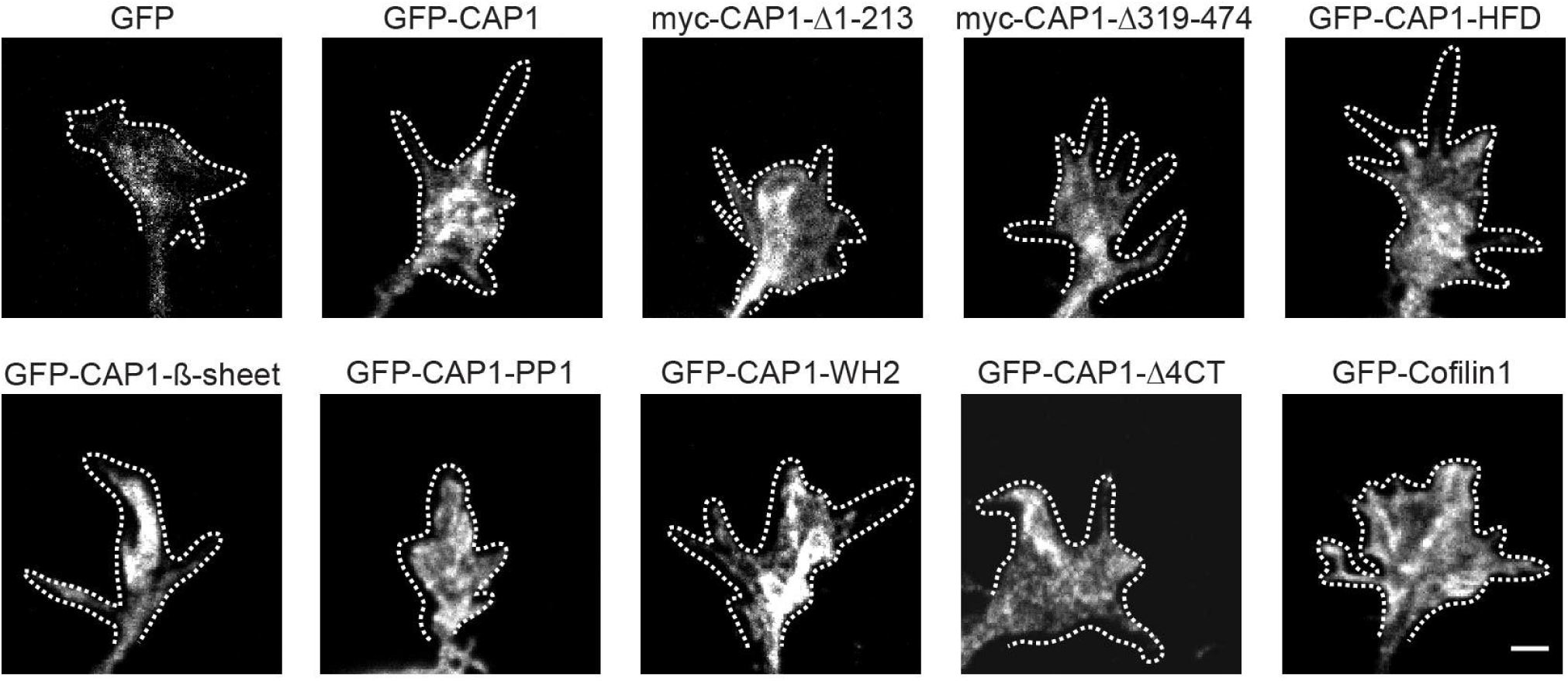
Localization of GFP, GFP-CAP1, mutant GFP-CAP1 variants, myc-CAP1 deletion constructs and GFP-cofilin1 in DIV1 growth cones. Dashed lines outline growth cones. Scale bars (in µm): 2.

**Figure S4.**
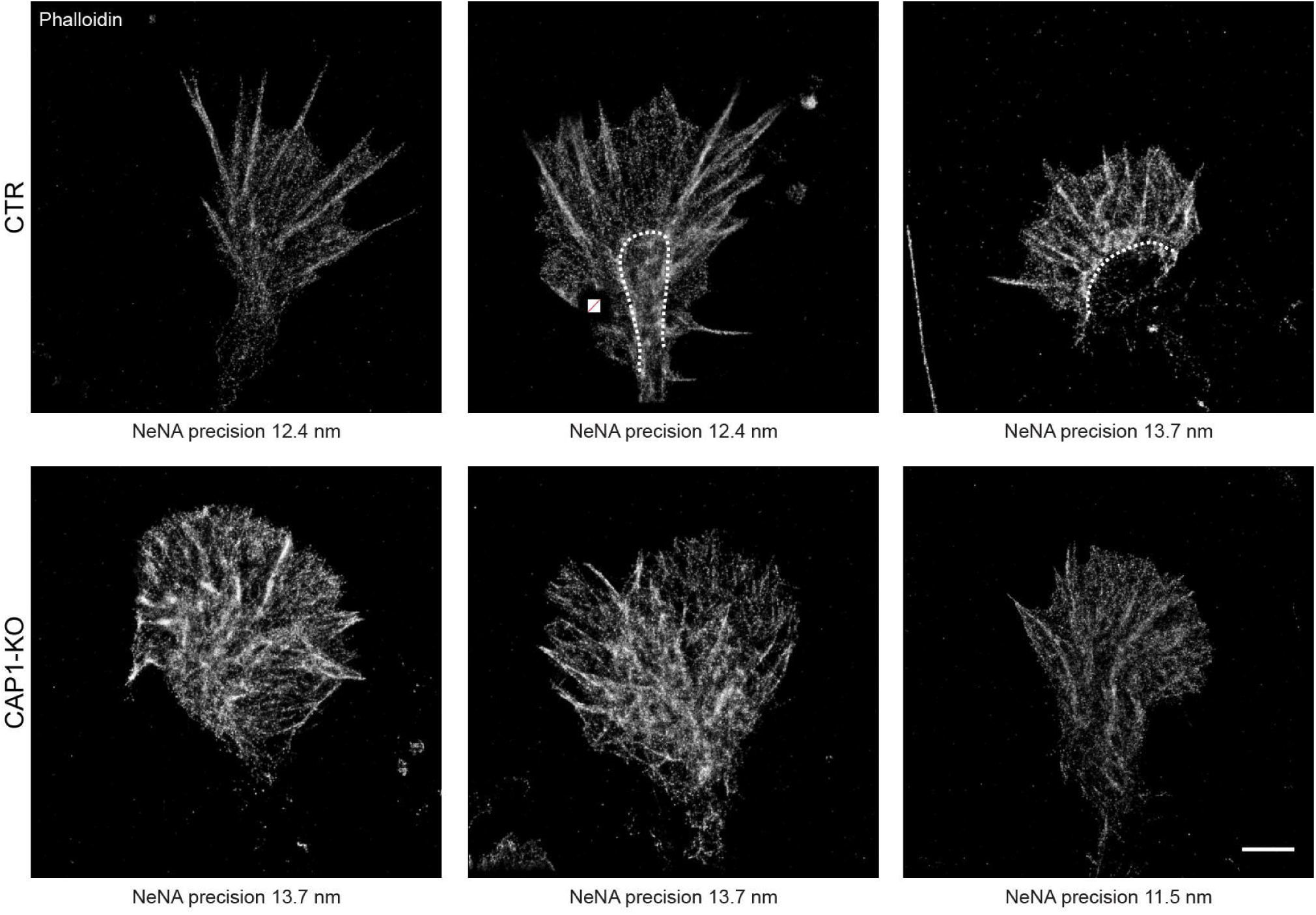
Micrographs of phalloidin-stained growth cones from CTR and CAP1-KO neurons acquired by dSTORM. Dashed line marks border between C domain and T zone. Crossed out box marks position of fiducial marker used for drift correction. Scale bar (in µm): 2.

**Figure S5.**
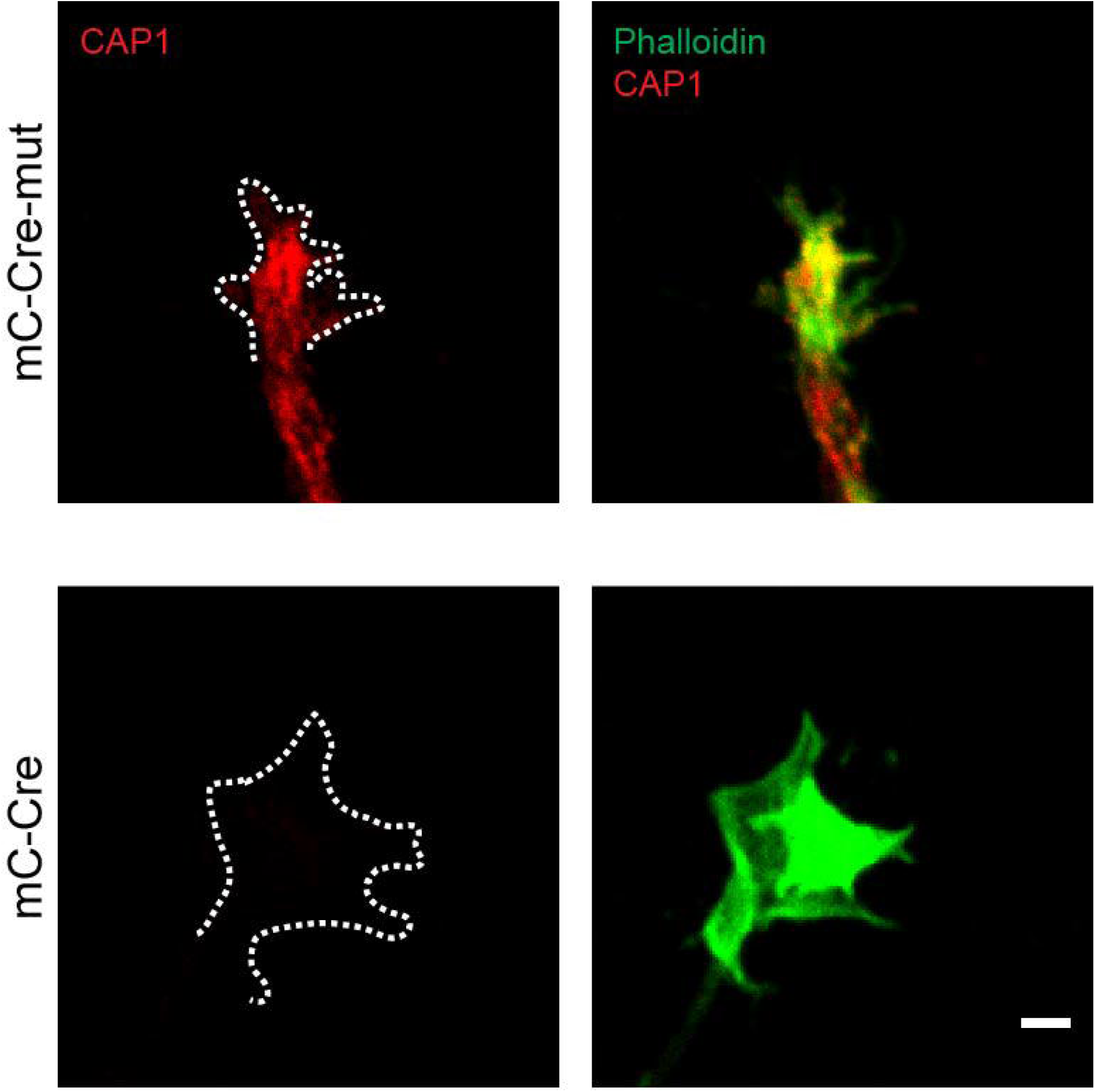
Micrographs of phalloidin and CAP1-stained growth cones from CAP1^flx/flx^/Cfl1^flx/flx^ growth cones either expressing mC-Cre-mut or mC-Cre. Dashed lines outline growth cones. Scale bar (in µm): 2.

**Movie S1:** Movie showing a CTR growth cone imaged by differential interference contrast (DIC) microscopy. Many protruding and retracting filopodia were present during image acquisition of 10 min. Scale bar: 2 µm.

**Movie S2:** Absence of filopodia and only low motility in a representative DIC imaged CAP1-KO growth cone. Acquisition time: 10 min, scale bar: 2 µm.

**Movie S3:** Movie showing GFP-actin recovery upon bleaching in a CTR growth cone. Upon bleaching fluorescence recovery was recorded over a time course of 3 min. Scale bar: 2 µm.

**Movie S4:** Movie showing GFP-actin recovery upon bleaching in a CAP1-KO growth cone. Upon bleaching fluorescence recovery was recorded over a time course of 3 min. Scale bar: 2 µm.

**Movie S5:** Movie showing a growth cone from a LifeAct-GFP-transfected CTR neuron. Images were acquired every 5 s for 5 min. Scale bar: 2 µm.

**Movie S6:** Movie showing a growth cone from a LifeAct-GFP-transfected CAP1-KO neuron. Images were acquired every 5 s for 5 min. Scale bar: 2 µm.

**Movie S7:** Movie showing GFP-actin recovery upon bleaching in a growth cone from a control neuron expressing catalytic inactive Cre (mC-Cre-mut). Upon bleaching fluorescence recovery was recorded over a time course of 3 min. Scale bar: 2 µm.

**Movie S8:** Movie showing GFP-actin recovery upon bleaching in a growth cone from a CAP1^flx/flx^ neuron expressing catalytic active Cre (mC-Cre). Upon bleaching fluorescence recovery was recorded over a time course of 3 min. Scale bar: 2 µm.

**Movie S9:** Movie showing GFP-actin recovery upon bleaching in a growth cone from a Cfl1^flx/flx^ neuron expressing catalytic active Cre (mC-Cre). Upon bleaching fluorescence recovery was recorded over a time course of 3 min. Scale bar: 2 µm.

**Movie S10:** Movie showing GFP-actin recovery upon bleaching in a growth cone from a CAP1^flx/flx^/Cfl1^flx/flx^ neuron expressing catalytic active Cre (mC-Cre). Upon bleaching fluorescence recovery was recorded over a time course of 3 min. Scale bar: 2 µm.

**Movie S11:** Movie showing a growth cone from a LifeAct-GFP-transfected control neuron expressing catalytic inactive Cre (mC-Cre-mut). Images were acquired every 5 s for 5 min. Scale bar: 2 µm.

**Movie S12:** Movie showing a growth cone from a LifeAct-GFP-transfected CAP1^flx/flx^ neuron expressing catalytic active Cre (mC-Cre). Images were acquired every 5 s for 5 min. Scale bar: 2 µm.

**Movie S13:** Movie showing a growth cone from a LifeAct-GFP-transfected Cfl1^flx/flx^ neuron expressing catalytic active Cre (mC-Cre). Images were acquired every 5 s for 5 min. Scale bar: 2 µm.

**Movie S14:** Movie showing a growth cone from a LifeAct-GFP-transfected CAP1^flx/flx^/Cfl1^flx/flx^ neuron expressing catalytic active Cre (mC-Cre). Images were acquired every 5 s for 5 min. Scale bar: 2 µm.

## Notes

### Competing Interest Statement

The authors have declared no competing interest.

